# Unprocessed serum glycosylphosphatidylinositol-anchored proteins are correlated to metabolic states

**DOI:** 10.1101/481549

**Authors:** Günter A. Müller, Andreas W. Herling, Kerstin Stemmer, Andreas Lechner, Matthias H. Tschöp

## Abstract

To study the possibility that components of eukaryotic plasma membranes are released in spontaneous or controlled fashion, a chip-based sensor was developed for complete glycosylphosphatidylinositol-anchored proteins (GPI-AP), which may form together with (phospho)lipids so far unknown (non-vesicular) extracellular complexes (GLEC). The sensor relies on changes in phase shift and amplitude of surface acoustic waves propagating over the chip surface upon specific capturing of the GPI-AP and detection of associated phospholipids and renders isolation of the labile GLEC unnecessary. GLEC were found to be released from isolated rat adipocyte plasma membranes immobilized on the chip, dependent on the flow rate and composition of the buffer stream. Moreover, incubation medium of isolated adipocytes and serum of rats are sources for GLEC which enables their differentiation according to cell size and genotype or body weight, respectively, as well as human serum.

---

<u>G</u>lycosyl<u>p</u>hosphatidylinositol-<u>a</u>nchored <u>p</u>roteins (GPI-AP), which represent about 1% of all proteins in eukaryotes, are constituted by a highly conserved hydrophobic glycolipidic membrane anchor (GPI) and variable large hydrophilic protein moieties^1-3^. On basis of their amphiphilic nature, <u>G</u>PI-AP equipped with the complete GPI anchor together with exogenous lipids (phospholipids, cholesterol), presumably required to shield their fatty acyl moieties from the aqueous environment, into <u>e</u>xtracellular <u>c</u>omplexes (GLEC) may be regarded as candidates for release from the extracellular face of plasma membranes upon exposure towards endogenous or exogenous cues, such as metabolites and mechanical forces. The putative release of GLEC was first studied with adipocytes since their plasma membranes undergo extensive stretching upon lipid filling and are in intimate contact with serum albumin and fatty acids. To avoid isolation of the presumably labile GLEC, a chip-based sensor was developed. It relies on specific capturing of the GLEC streaming through the microfluidic channels of the chip by their gold surface coated with α-toxin. Coating with α-toxin, which binds to the glycan core of the GPI anchor^4^, was performed with conventional coupling chemistry (**Supplementary Fig. 1**). Any (covalent or secondary) interaction of materials with the chip surface will lead to right-ward shifts in phase and/or reductions in amplitude of the horizontal <u>s</u>urface <u>a</u>coustic <u>w</u>aves (SAW) propagating along the chip surface. This reflects mass loading and/or increased viscosity, respectively, exerted by the interacting materials^5-7^. Consequently, the coating with α-toxin *per se* (**Supplementary Fig. 1**) and the capturing of GLEC can be monitored by chip-based sensing (see below).

**Figure 1.**
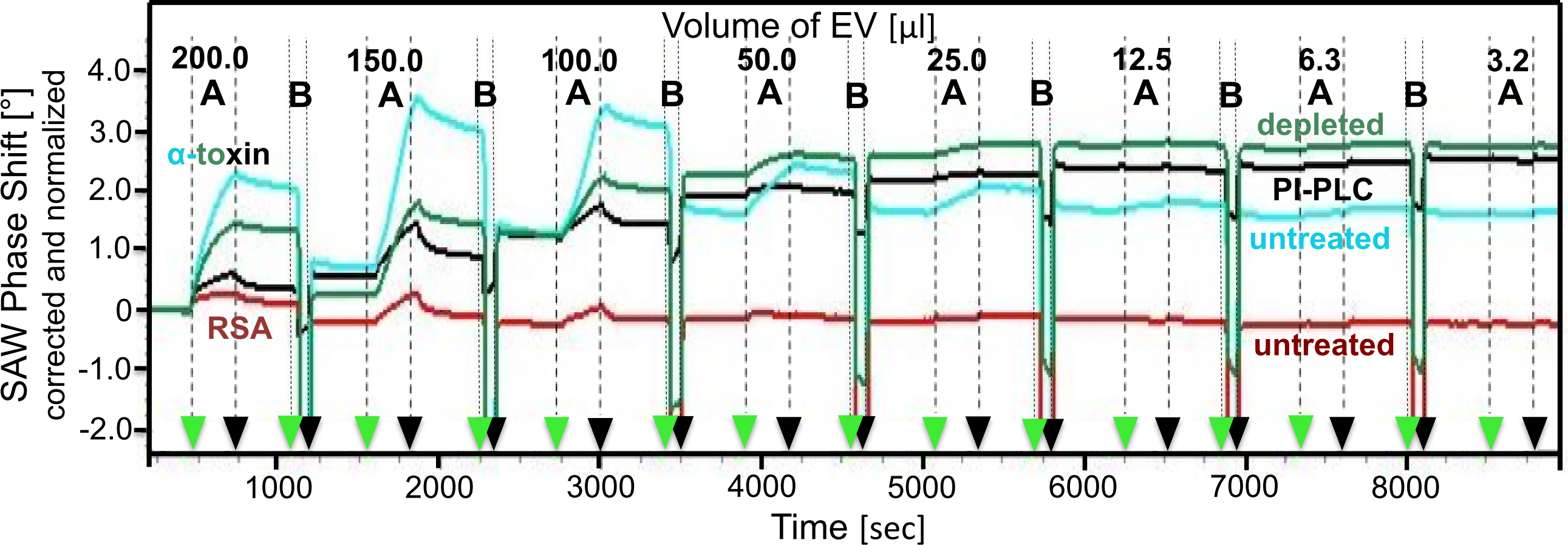
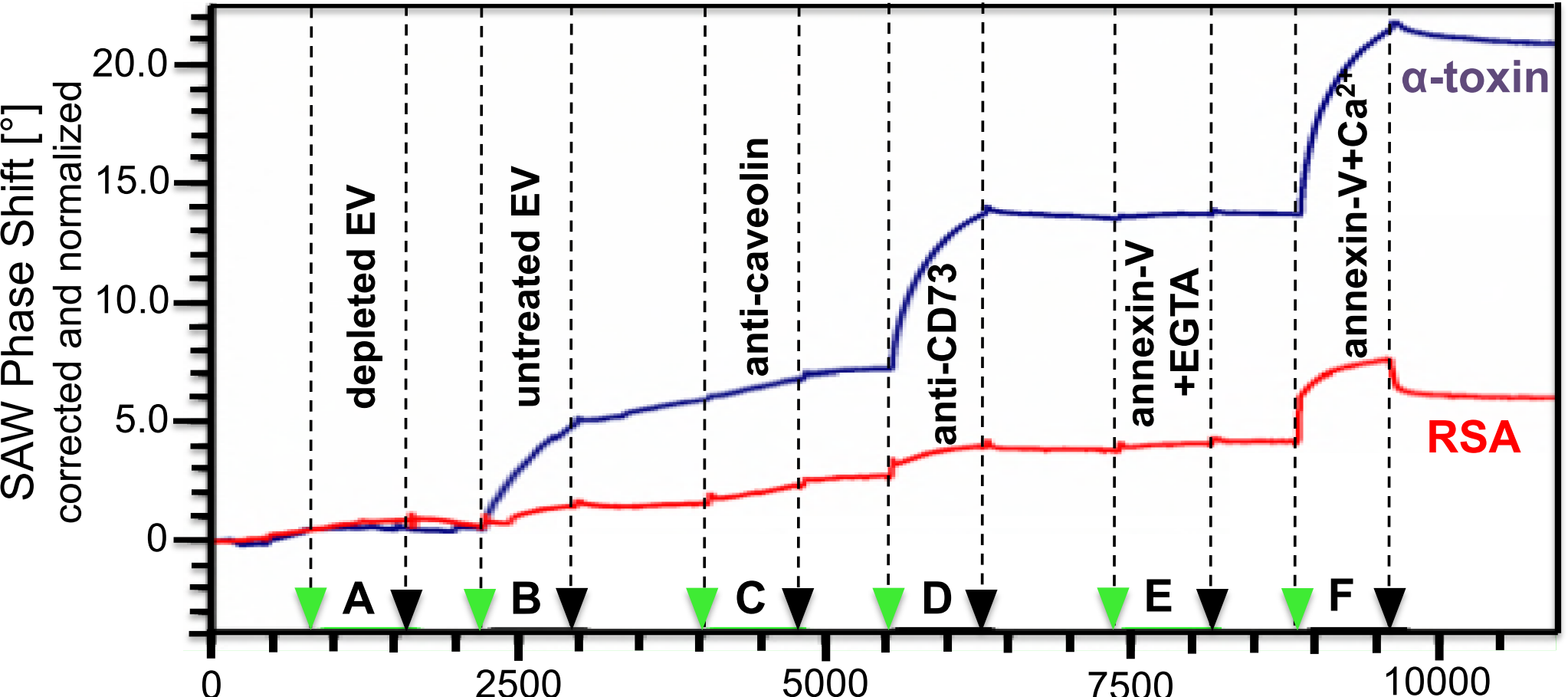
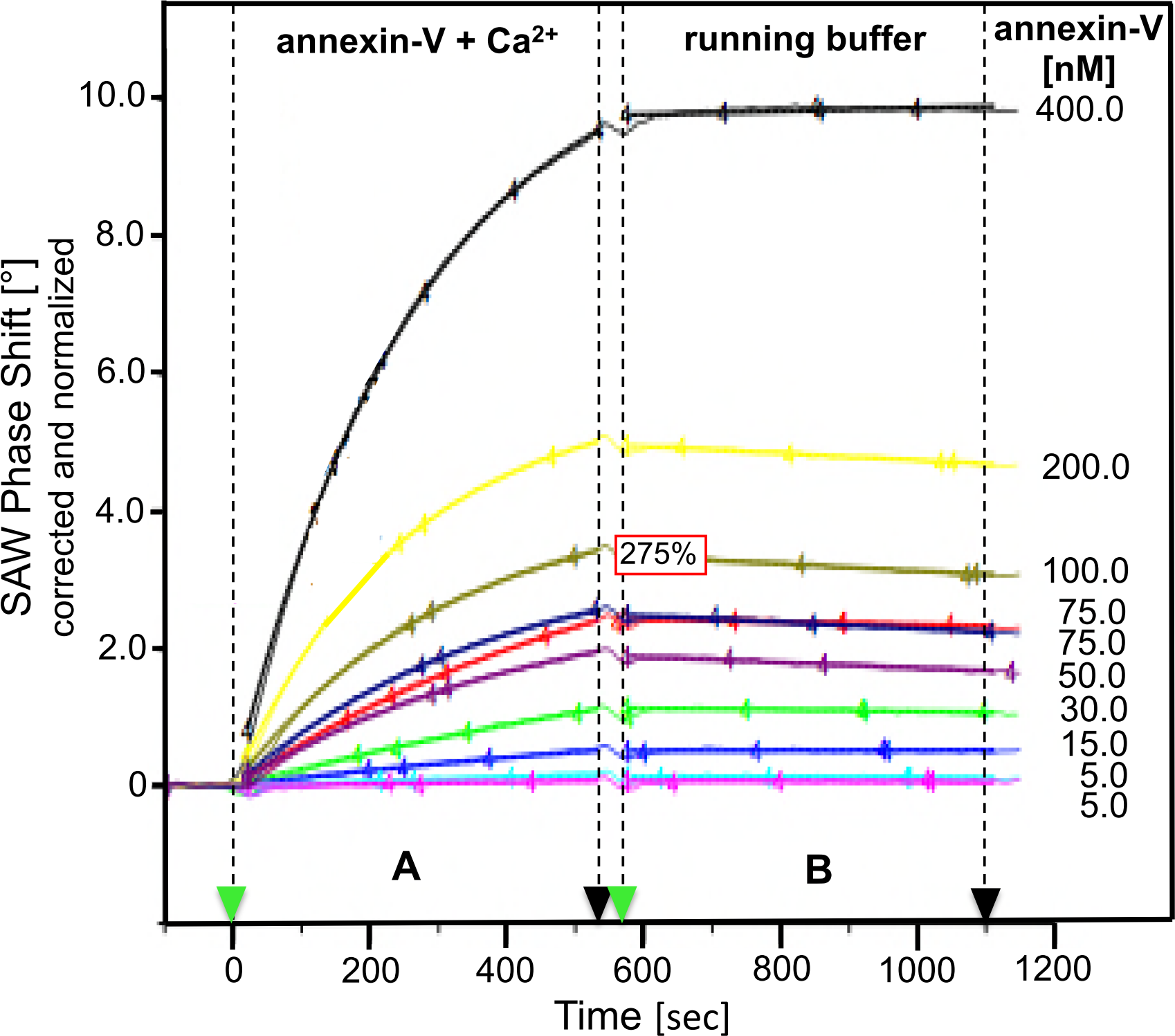
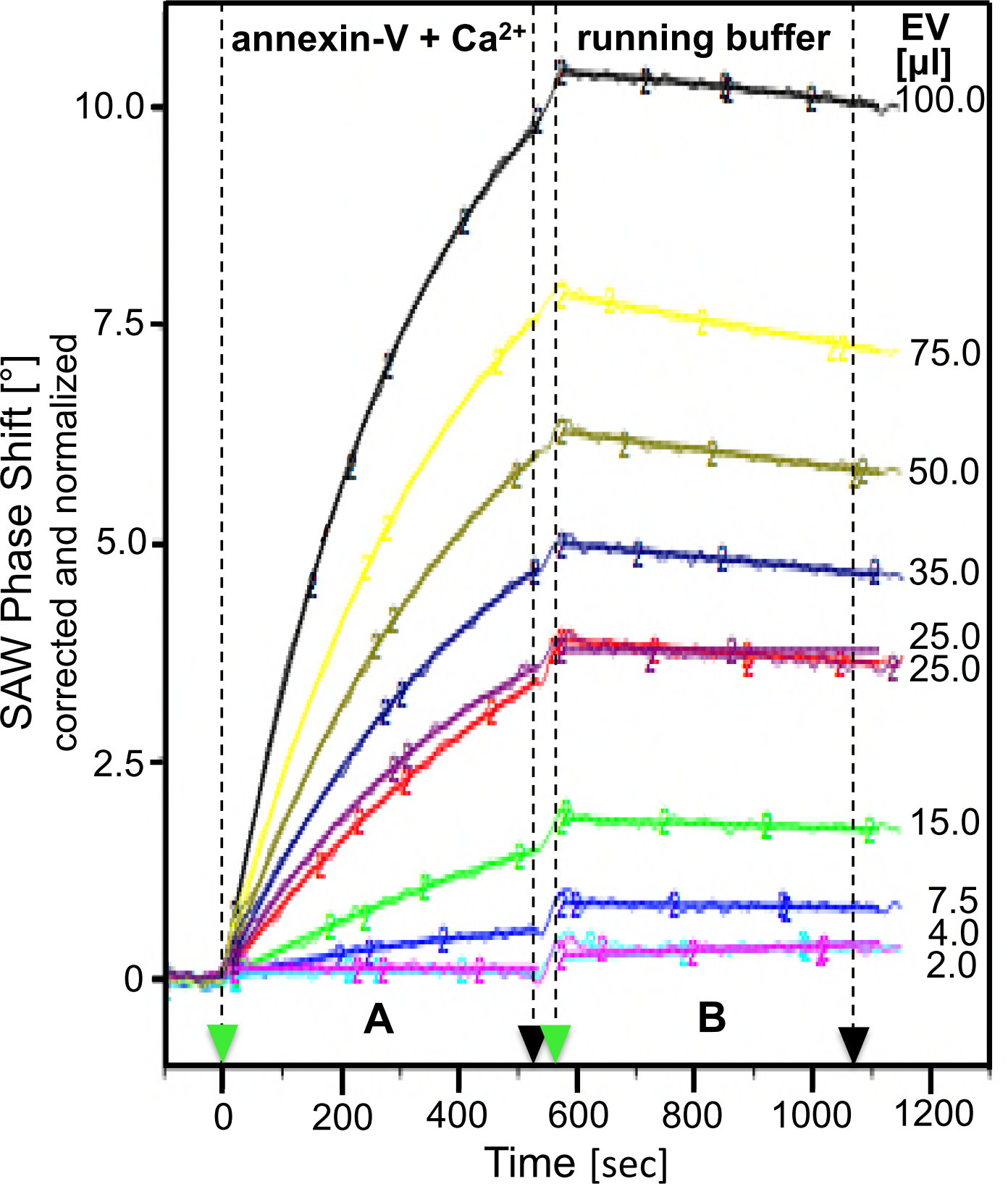
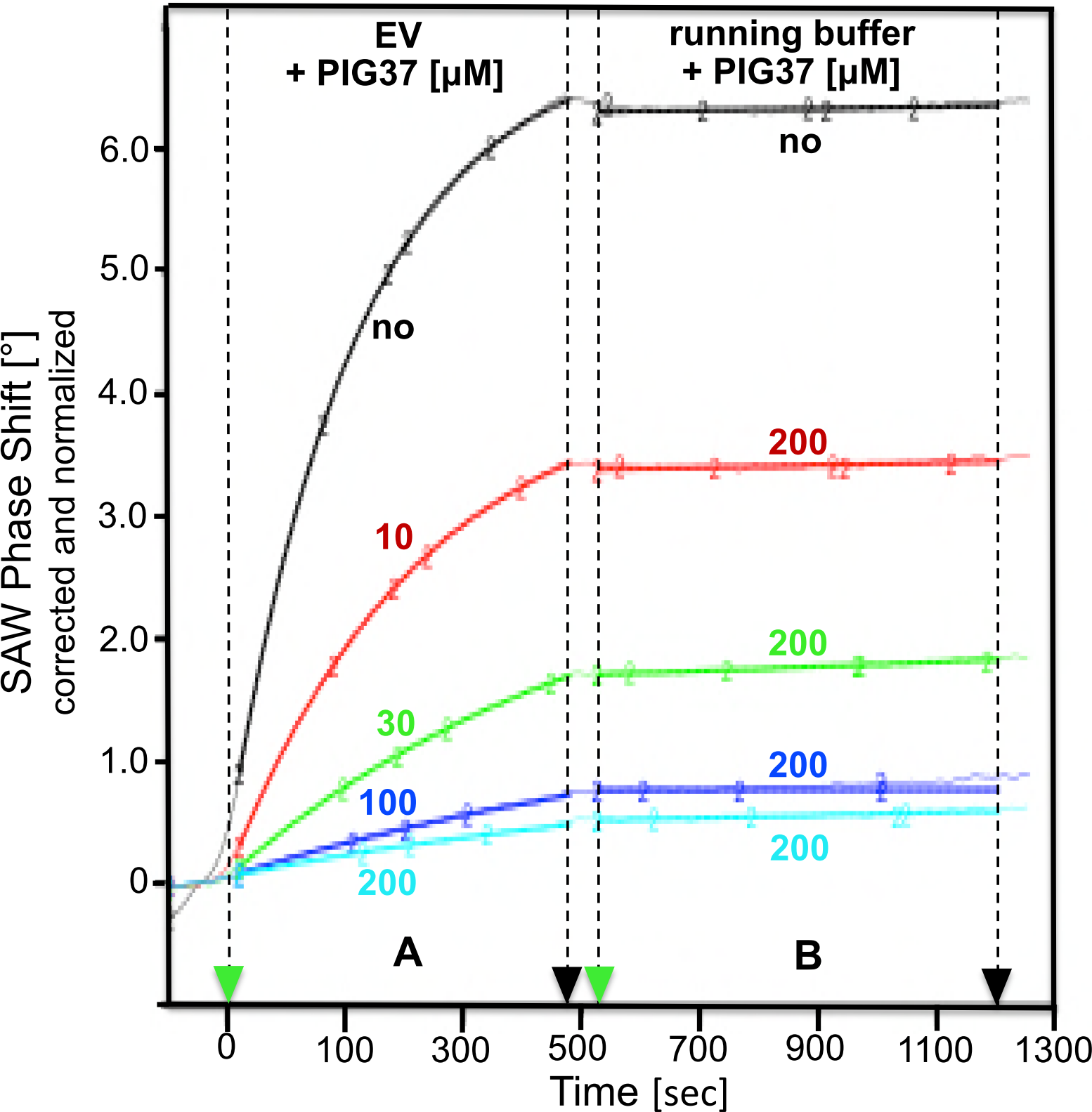
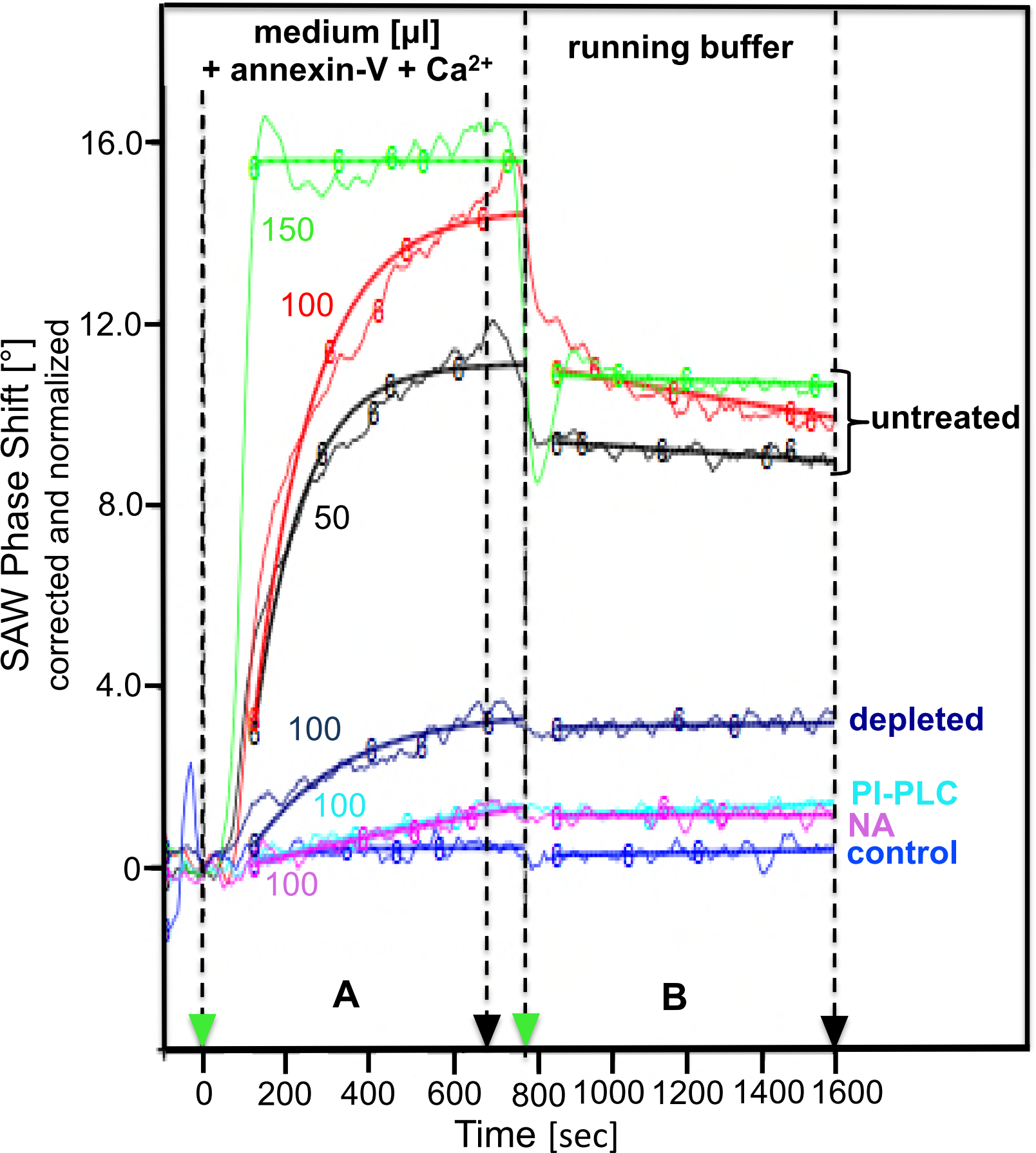
Implementation of the SAW chip-based sensing of GLEC. (**a**) For capturing of rat adipocyte EV, decreasing volumes of EV in EV buffer as indicated and adjusted to 200 μl final injection volume, which had remained untreated (turquoise and red curves) or had been pretreated with PI-PLC (black line) or had been depleted for GPI-AP (green curve), were sequentially injected into α-toxin-coated channels (turquoise, black and green curves) or RSA (red curve) or into “blank” uncoated channels (see below; period A). Phase shift is measured at a flow rate of 50 μl/min using EV buffer as running buffer and with regeneration after each injection (period B) and given (as °) upon correction for the “blank” channel and normalization (set at 0 for 250 sec; see **Supplementary Fig. 1a**). Representative diagram from four independent coating reactions is shown performed with distinct chips. (**b**) For “sandwich” detection of rat adipocyte EV by annexin-V and anti- CD73 antibodies, 162.5 μl of EV in EV buffer, which had been depleted for GPI-AP (period A) or left untreated (period B), were injected (at a flow rate of 13 μl/min) consecutively into α-toxin coated channels with (blue curve). For detection of unspecific binding of EV to serum albumin, 162.5 μl of 1% RSA in EV buffer were injected to yield the “albumin” channel by mere (non-covalent) adsorption (red curve). In addition, uncoated channels were run as “blank” channels. Subsequently, 162.5 μl each of 200 nM anti-caveolin antibodies (period C), 200 nM anti-CD73 antibodies (period D), 400 nM annexin-V containing 0.2 mM EGTA (period E) and 400 nM annexin-V containing 40 μM Ca^2+^ (period F) were injected successively into the channels. Phase shift is measured at a flow rate of 13 μl/min using EV buffer as running buffer and given (as °) upon correction for the “blank” channel (see **a**) and normalization (set at 0 for 0 sec). Representative diagram from five independent detection reactions is shown performed with three distinct chips and two distinct samples. A considerable phase shift was observed with RSA in the presence of annexin-V and Ca^2+^. Since albumin is a major component of serum and cell media, an “albumin” channel with RSA (in PBS) injected was included in the following experiments for correction of α-toxin-independent binding of annexin-V to the chip (possibly caused by its interaction with phospholipids associated with chip surface-adhering albumin from the serum or medium samples. (**c**) For studying the concentration-dependence of the “sandwich” detection of captured EV by annexin-V, 75 μl of EV in EV buffer were injected into α-toxin-coated channels or into a “blank” channel. Subsequently, 2×130 μl of annexin- V at the concentrations indicated in EV buffer containing 40 μM Ca^2+^ (period A) and then 2×130 μl of EV buffer as running buffer (period B) were injected into the channels at a flow rate of 30 μl/min. Phase shift is given (as °) upon correction for the “blank” channel and normalization (set at 0 for 0 sec and each annexin-V concentration) as both original and fitted data. Representative diagram from three independent detection reactions is shown performed with five distinct chips and the same sample. (**d**) For studying the volume- dependence of capturing and detection of adipocyte EV, increasing volumes of EV in EV buffer, adjusted to a total sample volume of 2×130 μl each, were injected into α-toxin- coated channels or into a “blank” channel. Subsequently, 2×130 μl of 400 nM annexin-V and 40 μM Ca^2+^ in EV buffer were injected into all channels (period A), followed by injection of EV buffer as running buffer at a flow rate of 30 μl/min (period B). Phase shift is given (as °) after correction for the “blank” channel and normalization (set at 0 for 0 sec and each volume of EV) as both original and fitted data. Representative diagram from three independent detection reactions is shown performed with six distinct chips and two distinct samples. (**e**) For studying the effect of PIG37 on capturing of EV, 150 μl of EV in EV buffer in the absence (black curve) or presence of PIG37 at the concentrations indicated (red, green, pink, blue curves) were injected into α-toxin-coated channels or into a “black” channel at a flow rate of 20 μl/min (period A). Subsequently, 225 μl of EV buffer containing 200 μM PIG37 or lacking PIG37 was injected at a flow rate of 20 μl/min (period B). Phase shift is given (as °) upon correction for the “blank” channel and normalization (set at 0 for 0 sec and 200 μM PIG37; blue curve) as original and fitted data. Representative diagram from two independent detection reactions is shown performed with two distinct chips and two distinct samples. (**f**) For detection of GLEC in adipocyte incubation medium, increasing volumes of medium obtained by incubation of rat adipocytes of medium size and kept untreated (black, red, green curves) or depleted for GPI-AP (dark blue curve) or pretreated with PI-PLC (light blue curve) or nitrous acid (NA; pink curve) or medium lacking adipocytes (control; violet curve) and adjusted to 150 μl total volume with adipocyte buffer each were injected into α-toxin-coated channels. Subsequently, 2×150 μl of 20 mM TRIS/HCl (pH 8.0), 150 mM NaCl, 250 mM sucrose containing 400 nM annexin-V and 40 μM Ca^2+^ were injected at a flow rate of 25 μl/min (period A). Thereafter, 1 ml of 20 mM TRIS/HCl (pH 8.0), 150 mM NaCl and 250 mM sucrose were injected at a flow rate of 75 μl/min (period B). Phase shift is given (as °) upon correction for the “blank” channels and normalization (set at 0 for 0 sec and 150 μl volume) as original and fitted data. Representative diagram from three independent detection reactions is shown with the same chip and two distinct (medium) samples.

Initially, the sensor was developed and validated using so-called extracellular vesicles (EV) as analytes. EV are membrane vesicles which are released from most cell types^8^, in particular upon challenge with exogenous stressors^9^. A subset of EV released from adipocytes into the incubation medium are known to harbor complete GPI-AP at the outer surface of their phospholipid bilayer^10,11^ and consequently can be regarded as subtype of GLEC. Injection of EV isolated from rat adipocyte incubation medium into α-toxin-coated, but not albumin-coated chips caused volume-dependent phases shifts of the SAW (**Fig. 1a**). Depletion of the GPI-AP-harboring EV upon adsorption to α-toxin-coupled magnetic beads or cleavage of the GPI anchor by bacterial PI-PLC prior to injection (partially) prevented phase shift. The presence of GPI-AP, such as CD73, and phospholipids in the EV was shown by sequential binding “in sandwich” of anti-CD73 antibodies and the Ca^2+^- dependent phospholipid-sequestering protein annexin-V (in the presence of Ca^2+^, but not EGTA) to the chip (**Fig. 1b**). The specificity of detection of GPI-AP in association with lipids was confirmed by the lack of SAW phase shift using (i) chips (non-covalently) coated with rat serum albumin, (ii) EV depleted from GPI-AP or (iii) antibodies against the membrane protein caveolin, located at the luminal membrane leaflet of the EV (**Fig. 1b**) as well as by the dependence of SAW phase shift on the annexin-V concentration (**Fig. 1c**) and sample volume (**Fig. 1d**) during period A. The specificity of capturing of the EV as GLEC subtype *via* α-toxin was confirmed by maintenance and concentration-dependent reduction of phase shift during extensive washing of the chip surface with running buffer (**Fig. 1c-f,** periods B) and presence of synthetic <u>p</u>hosphoinositol<u>g</u>lycan (PIG) 37 mimicking the glycan core moiety of GPI-AP^12^ during the association period (**Fig. 1e**, period A), respectively. Subsequent washing of the captured EV with PIG37 at high concentration did not cause phase shift decrease, as was true for running buffer alone, reflecting a PIG37- dependent delay in association rather than induction of dissociation (**Fig. 1e**). This is presumably due to the structural deviation of PIG37 from the authentic GPI glycan core, since PIG41 resembling it more closely^12^ managed the concentration-dependent displacement of captured EV as well as blockade of re-capturing of the displaced EV (**Supplementary Fig. 2**).

**Figure 2.**
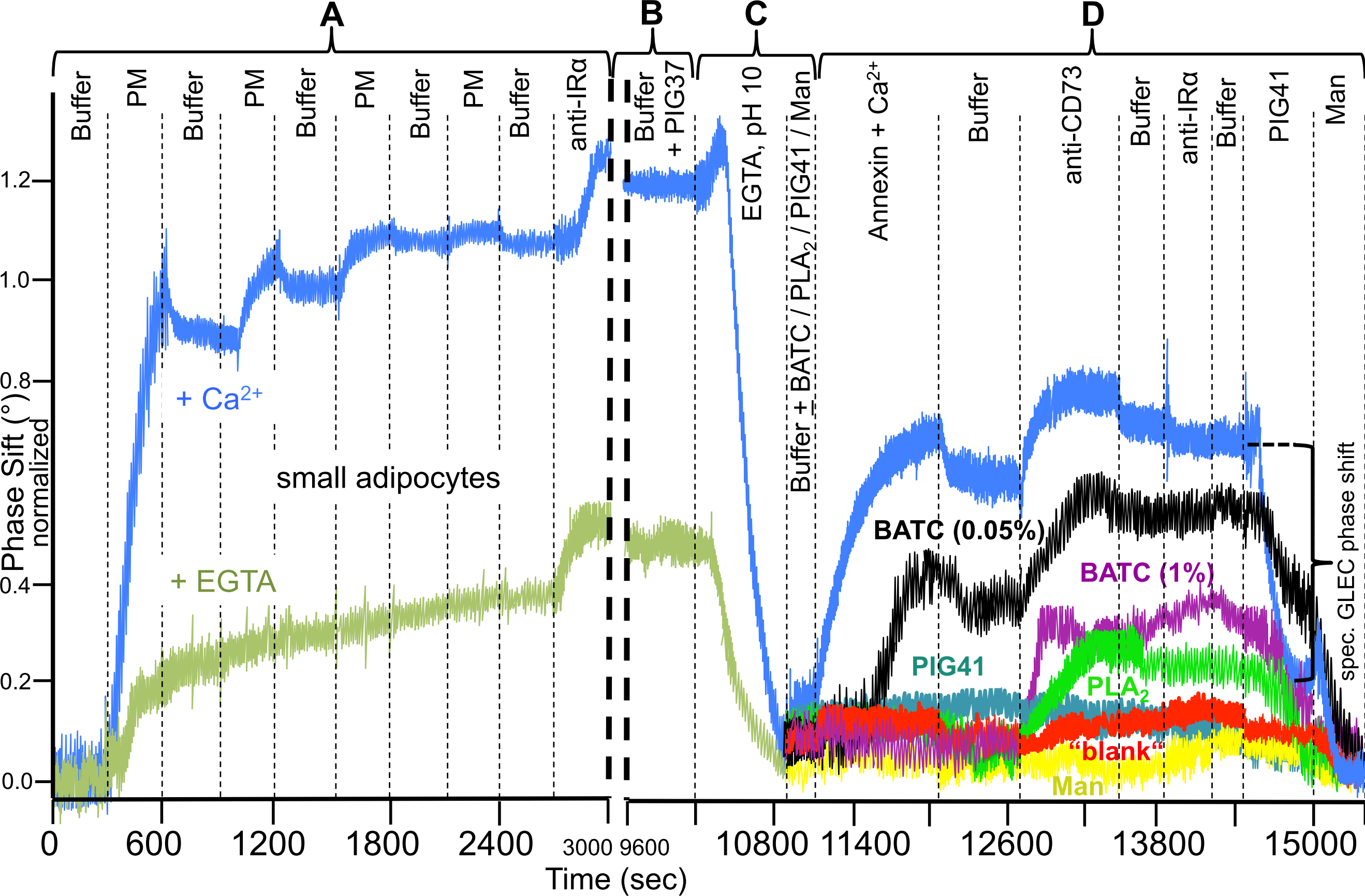
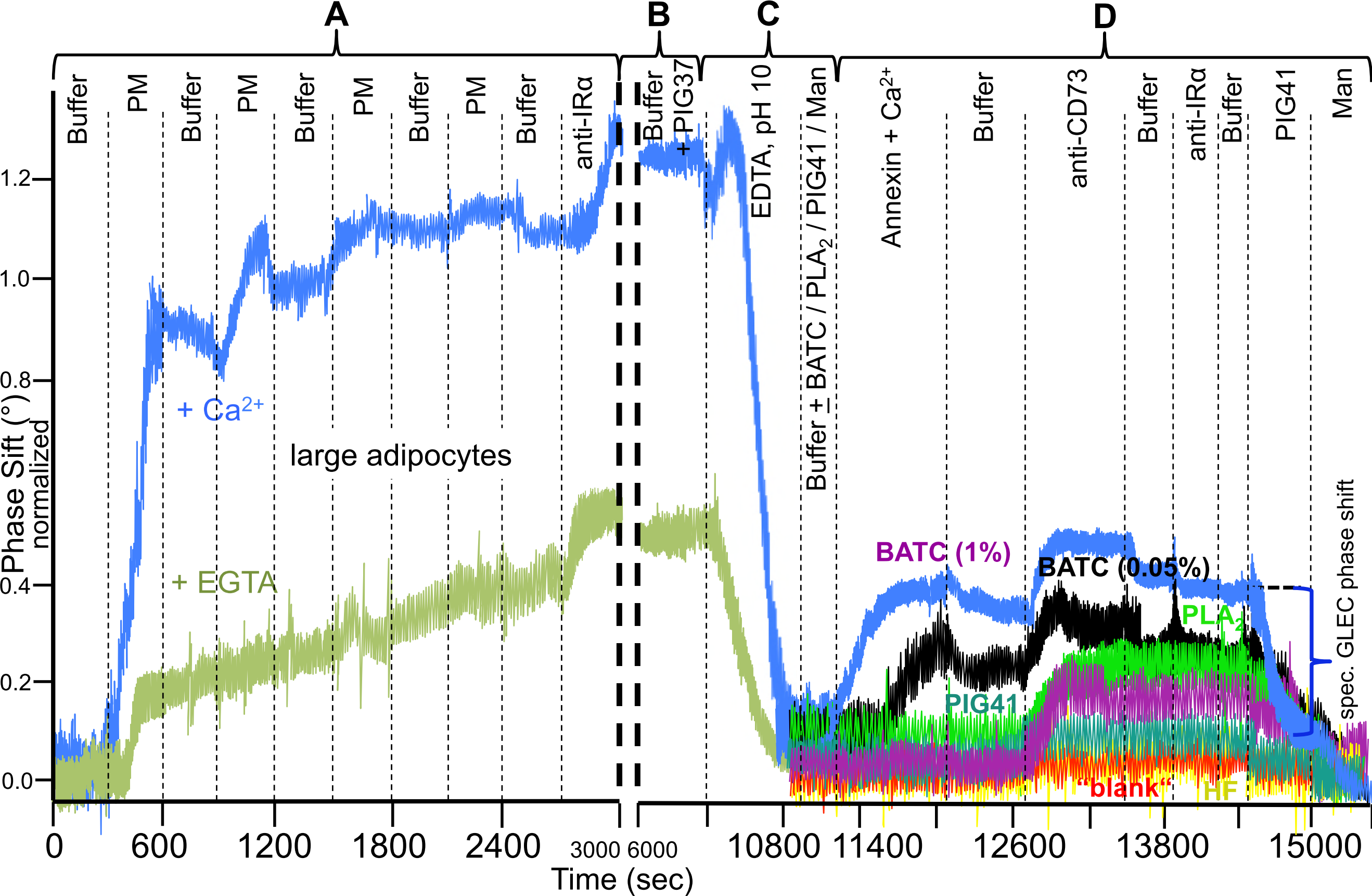
Implementation of the “lab-on-the-chip”-based sensing of plasma membrane-derived GLEC. For immobilization of plasma membranes from small. (**a**) or large (**b**) adipocytes, three 100-μl portions in 10 mM HEPES/KOH (pH 7.5), 150 mM NaCl and 100 mM sucrose (PM) in the presence of 2 mM EGTA (light green curve) or 1 mM Ca^2+^ (blue curve) and, alternatively, three 100-μl portions of running buffer were consecutively injected into α-toxin-coated channels or uncoated “blank” channel (red curve) at a flow rate of 20 μl/min at 20°C (period A). For demonstration of the presence of the insulin receptor α-chain (IRα), 100 μl of 25 nM anti-IRα antibodies were injected at the end of period A. For the putative generation of GLEC from the immobilized plasma membranes and their capturing by the α-toxin-coated chip, 24 ml of running buffer containing 30 μM PIG37 were injected at a flow rate of 200 μl/min at 37°C (period B). For release of the plasma membranes from the chip surface, 1 ml of 10 mM glycine (pH 10) containing 2 mM EGTA were injected at a flow rate of 100 μl/min, followed by injection of 300 μl of running buffer alone (blue and red curves) or buffer containing 0.05% BATC (black curve), 1.5% BATC (pink curve), PLA_2_ (green curve), 200 μM PIG41 (turquoise curve) or 100 mM mannose (yellow curve) at a flow rate of 100 μl/min at 20°C (period C). For demonstration of capturing of GLEC, 200 μl of 400 nM annexin-V, 40 μM Ca^2+^, 10 mM HEPES/KOH (pH 7.5), 150 mM NaCl and 100 mM sucrose, 150 μl of running buffer, 180 μl of 200 nM anti-CD73 antibodies, 10 mM HEPES/KOH (pH 7.5), 150 mM NaCl and 100 mM sucrose, 90 μl of running buffer, 90 μl of 25 nM anti-IRα antibodies, 10 mM HEPES/KOH (pH 7.5), 100 mM sucrose, 150 mM NaCl and 0.2 mM EGTA, 65 μl of running buffer, 130 μl of 200 μM PIG41, 10 mM HEPES/KOH (pH 7.5), 100 mM sucrose, 150 mM NaCl and 0.2 mM EGTA and finally 90 μl of 10 mM HEPES/KOH (pH 7.5), 150 mM NaCl and 100 mM mannose were injected at a flow rate of 13 μl/min at 20°C in that sequential fashion (period D). Phase shift (as °) is given upon normalization (set at 0 for 0 sec). Representative overlay diagrams from six independent runs for each condition (periods A-D) are shown performed with three distinct chips (and the channels re-used six times), the same samples and the same instrument. The calculation of the specific GLEC phase shift as the difference between the total (normalized) phase shift after binding “in sandwich” of both annexin and anti-CD73 antibodies with subsequent washings (at 14500 sec) and the unspecific phase shift left after injection of PIG41 (1500 sec) is indicated.

Next, total incubation medium, which contains EV at rather low concentration only, was used to test the sensitivity and the potential to differentiate GLEC subtypes (putatively of non-vesicular configuration) between adipocytes of different size. Medium from primary rat adipocytes elicited volume-dependent (during capturing by α-toxin and detection by annexin-V, period A) and stable (during subsequent washing with running buffer, period B) phase shift increases, which were abrogated in course of depletion of the GPI-AP or cleavage of the GPI anchor by bacterial PI-PLC and nitrous acid deamination (**Fig. 1f**). Interestingly, media from large, medium and small adipocytes (**Supplementary Table 1)** elicited slightly different kinetics for capturing of the GLEC and subsequent detection of phospholipids (**Supplementary Fig. 3**, period A), which were both prevented by the detergent NP-40 (B). The injection of 10 and 200 μM PIG37 at the midst of capturing (**Supplementary Fig. 4a**, period A>B) led to maximal differentiation between GLEC from large *vs*. medium/small and large/medium *vs*. small adipocytes, respectively. 30 μM PIG37 turned out to enable the parallel discrimination of GLEC from large, medium and small cells considering phase shift (**b,** periods A1/2>B1/2) and amplitude reduction (**c**, period A2>B2), even in the presence of annexin-V during capturing of GLEC for simultaneous detection of phospholipids (period A2). In the absence of PIG37 minor differences between adipocytes of differing size were measured, only (period B1). Final injection of PIG41 caused complete abrogation of the medium-induced phase shift (**b**, period C1/2) as well as amplitude reduction (**c**, period D1/2). This total dissociation of GLEC from the chip argues for the specificity of their capturing and the possibility of their re-use. Importantly, two distinct chips run in parallel displayed very similar kinetics of capturing and dissociation (**b**). Taken together, the findings obtained with isolated EV and total incubation medium from rat adipocytes strongly suggest that chip-based sensing enables the sensitive identification of GLEC, i.e. EV and possibly non-vesicular subtypes, as well as their differentiation according to the releasing cell (here adipocytes of different size) on basis of the kinetics of the PIG37-dependent phase shift and amplitude reduction. In contrast, differentiation was not feasible on basis of the patterns of total proteins, total GPI-AP or specific GPI-AP contained in adipocyte incubation media (**Supplementary Fig. 5**).

**Figure 3.**
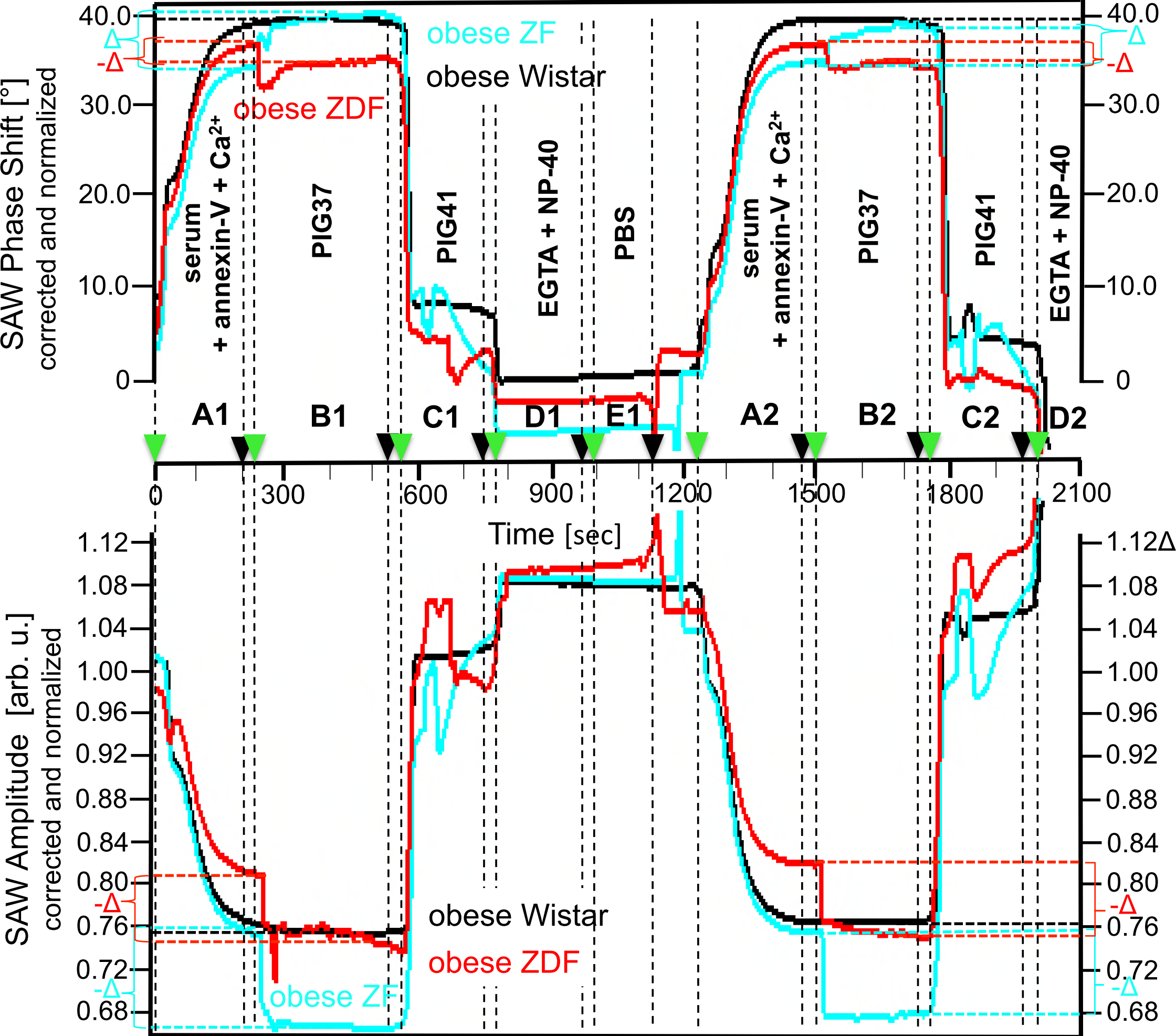
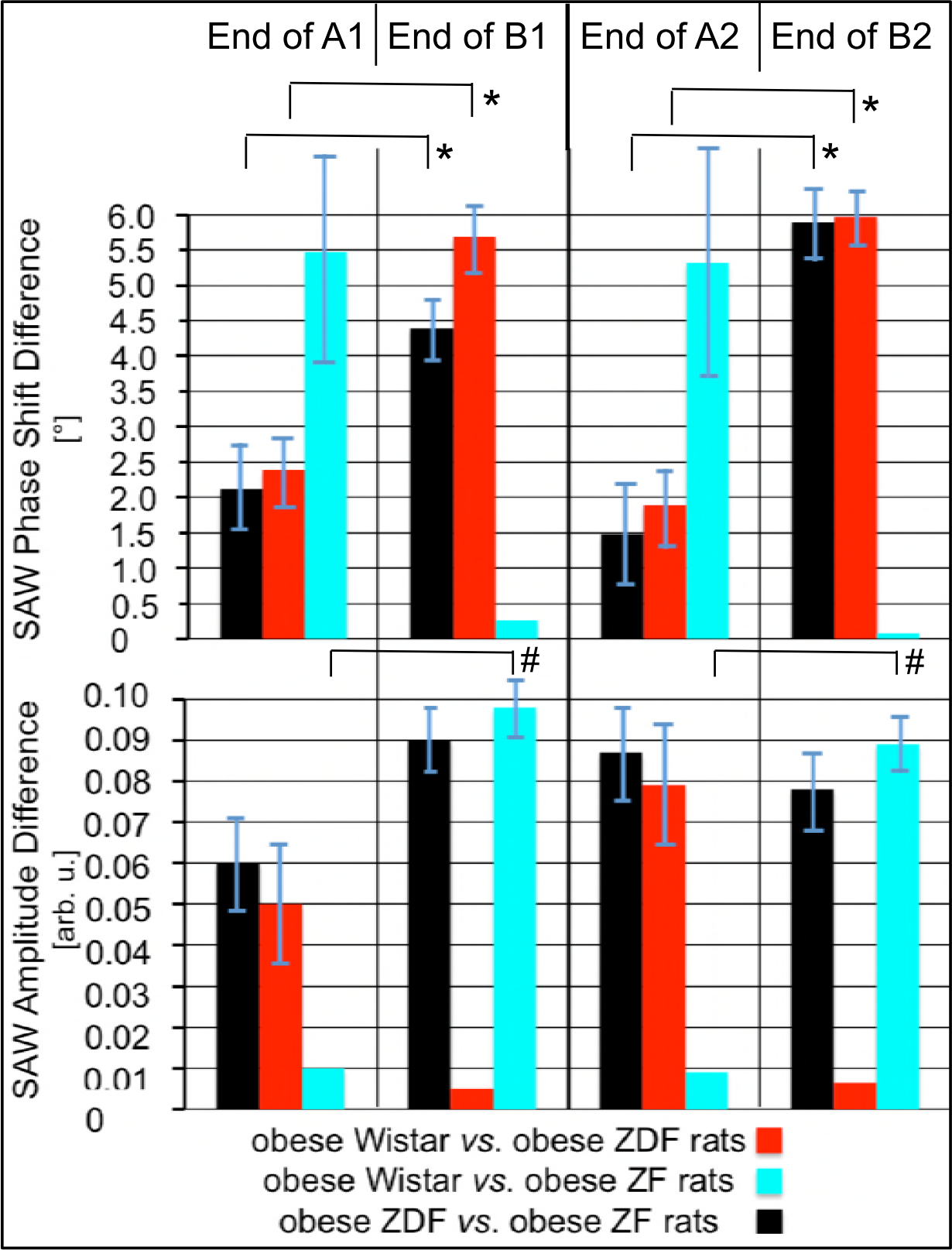
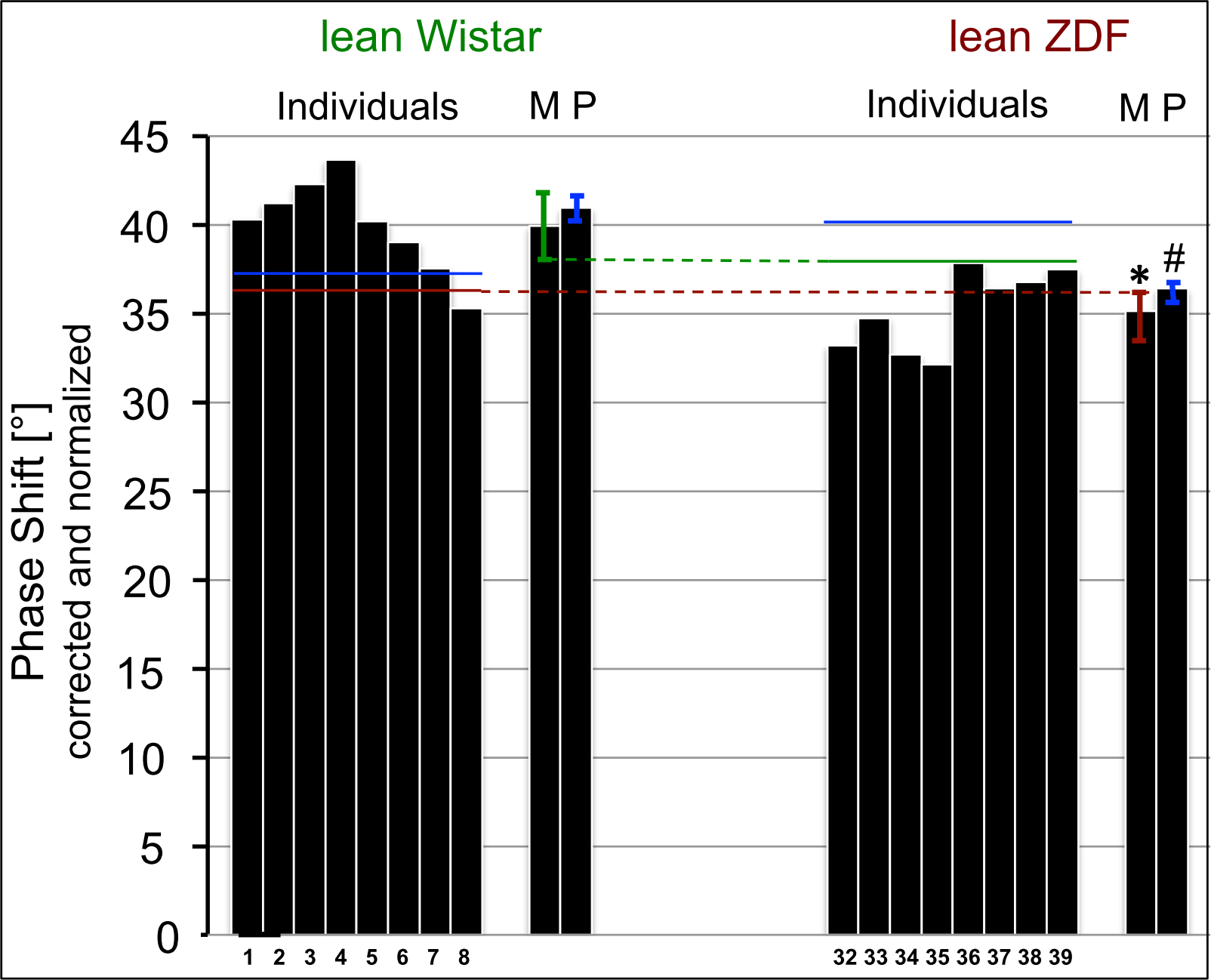
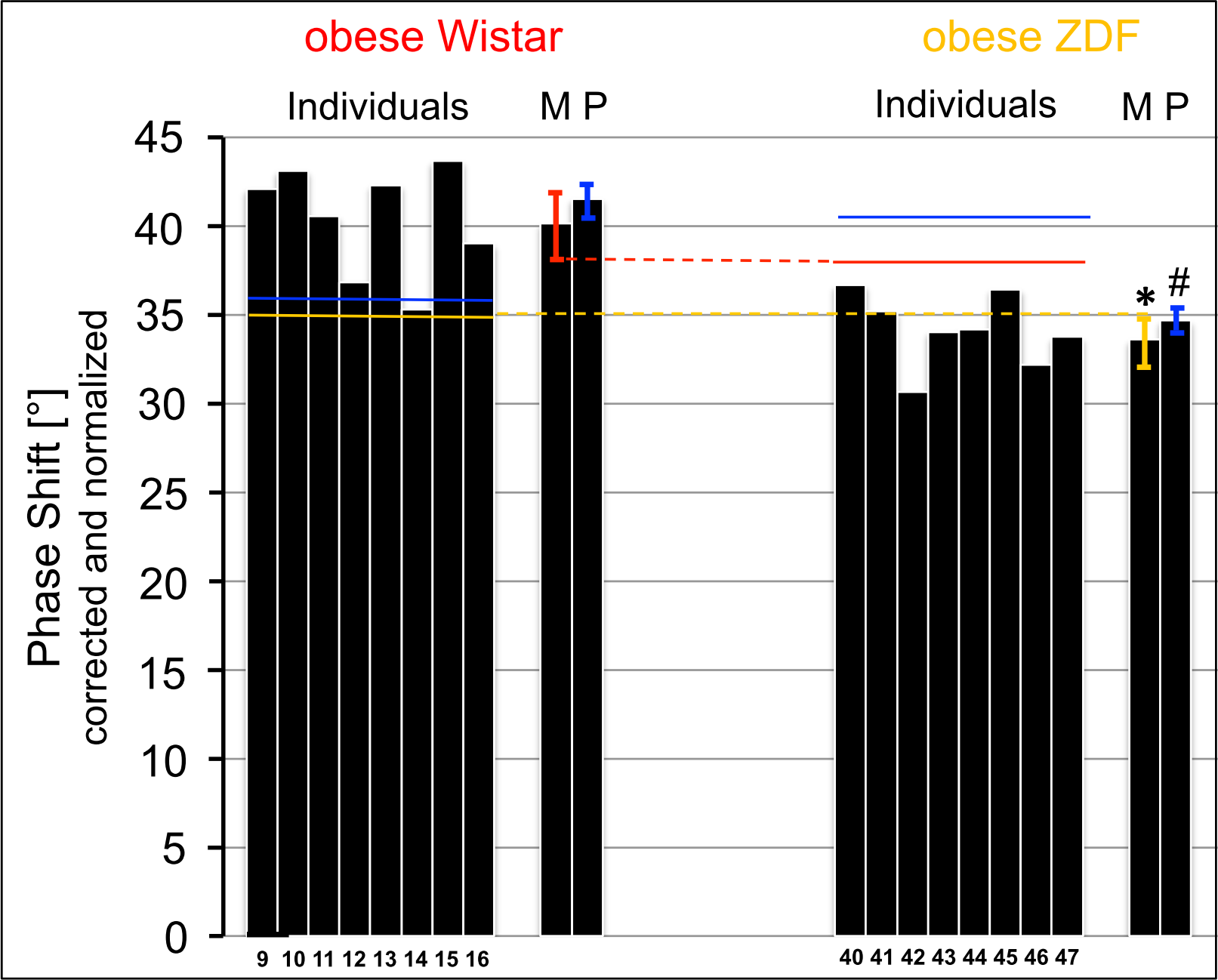
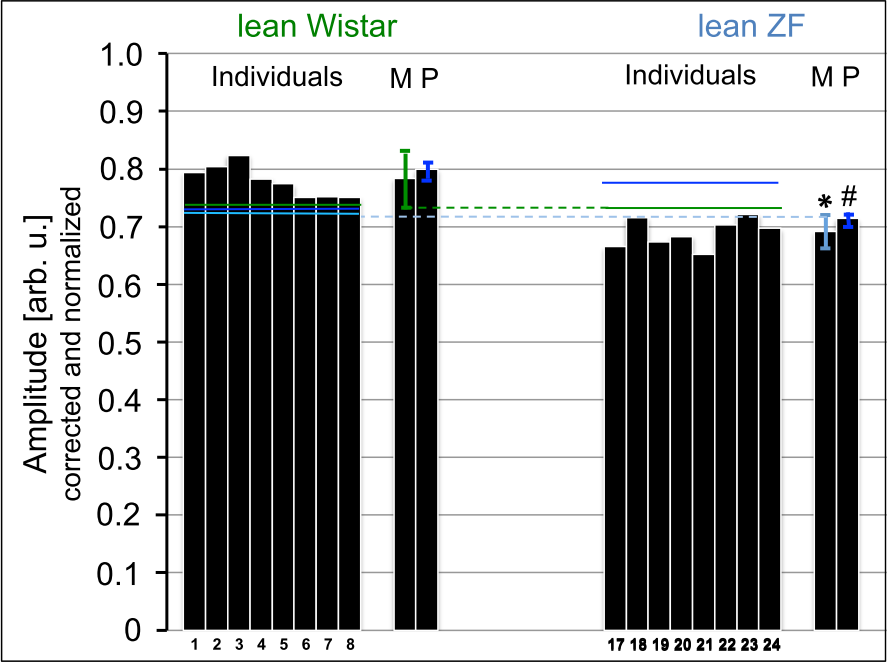
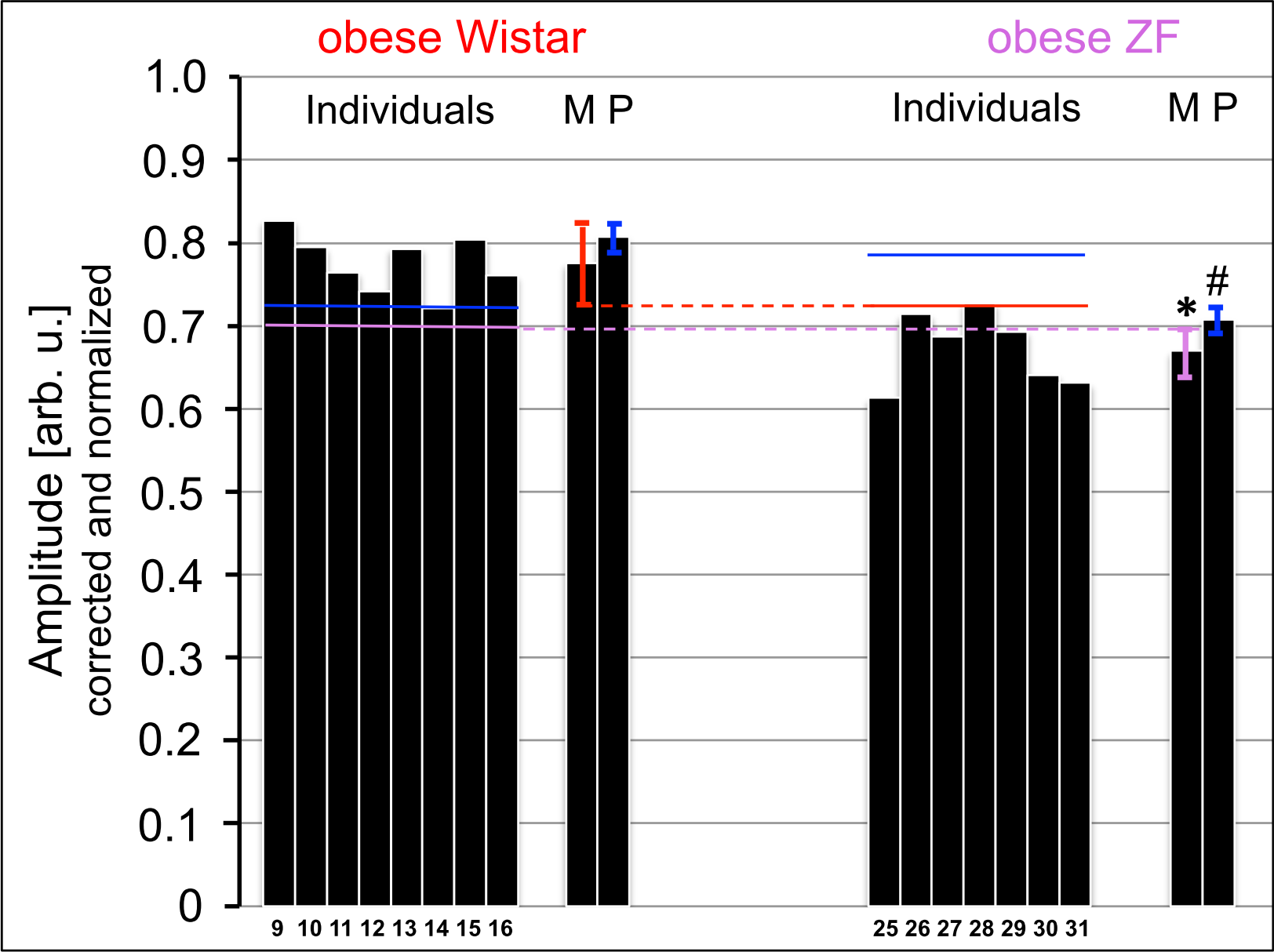
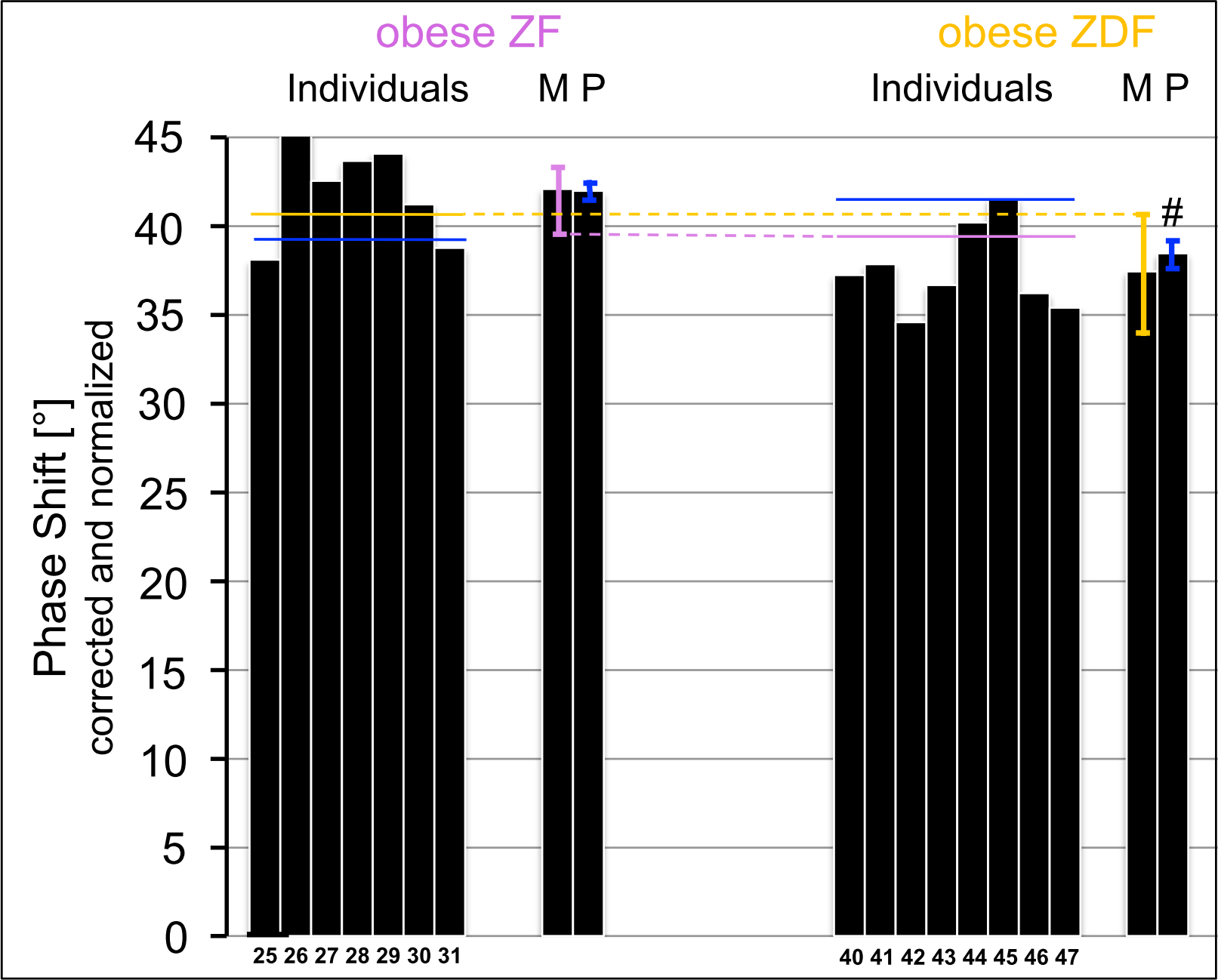
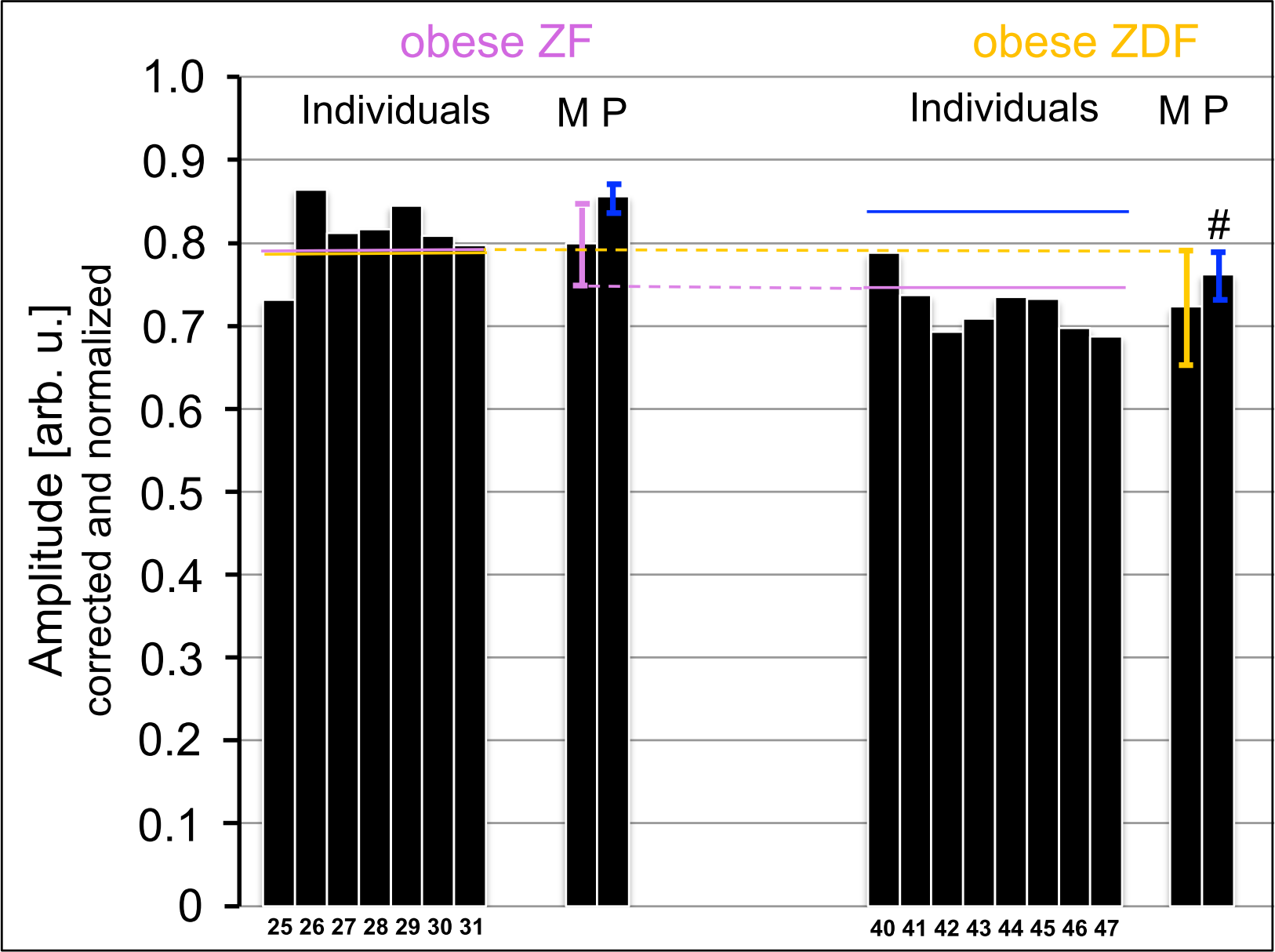
SAW chip-based sensing of GLEC in rat serum. (**a**) For PIG37-dependent differentiation between obese Wistar, ZF and ZDF rats upon simultaneous capturing and detection of GLEC, 40 μl of pooled serum sample from obese Wistar (black curve), ZF (turquoise curve) and ZDF (red curve) rats diluted five-fold with PBS and incubated (10 min, 20°C) with 200 μl of 800 nM annexin-V and 80 μM Ca^2+^ were injected into α-toxin- coated channels at a flow rate of 120 μl/min and at 20°C (period A1). To initiate the differentiation, 100 μl of 30 μM PIG37 in PBS were injected at a flow rate of 20 μl/min and at 37°C (period B1). To regenerate the chips, 50 μl of 200 μM PIG41 in PBS were injected at a flow rate of 15 μl/min for displacement of GLEC (period C1), followed by injection of 2×150 μl of PBS containing 0.2 mM EGTA and 0.05% NP-40 at a flow rate of 90 μl/min for removal of phospholipids (period D1) and finally of 2×100 μl of PBS at a flow rate of 90 μl/min (period E1). To demonstrate reproducibility of this protocol, the periods A-D were repeated under identical conditions with slightly adapted time frames. Phase shift (upper panel) and amplitude (lower panel) are given (in ° and arb. unit, respectively) after correction for the “blank” and “albumin” channels and normalization (set at 0° and 1 arb. unit, respectively, for start of the injection at 0 sec) as original data. The maximal phase shift and minimal amplitude, respectively, measured after the injection of PIG37 at the end of period B are indicated as hatched lines. The differences in maximal phase shift and minimal amplitude between obese Wistar and ZDF rats are indicated (red Δ), obese Wistar and ZF rats (turquoise Δ) and obese ZDF and ZF rats (black Δ) as measured at the end of period B1/2 (presence of PIG37) with Δ representing increases and -Δ decreases. Representative diagram from six independent runs for each serum type is shown performed with two distinct chips (re-used four times) and the same samples. (**b**) The differences (absolute values) in phase shift [°] and amplitude [arb. units] between obese Wistar and ZDF (red bars), obese Wistar and ZF (turquoise bars) and obese ZDF and ZF (black bars) rats measured at the end of period A1/2 (absence of PIG37) or period B1/2 (presence of PIG37) are given as means ± SD (* *p* ≤ 0.05; ^#^ *p* ≤ 0.01) calculated from four independent runs for each serum type performed with four distinct chips (re-used six times) and the same samples and instrument. In the following experiments the phase shift and amplitude were measured at the end of period B. (**c-h**) For the comparative analysis of serum GLEC from rats of similar body weight and different genotype, phase shift (**c**, **d**, **g**) and amplitude (**e**, **f**, **h**) were measured in the presence of 30 μM PIG37 as described for (**a**) are given for the individual samples and the means M ± SD calculated thereof as well as the pooled samples P ± SD derived from 12 (M) and 4 (P) independent runs for each sample performed with one chip (re-used six times) and 12/4 setting for each rat group (^#^ *p* ≤ 0.05, * *p* ≤ 0.01).

**Figure 4.**
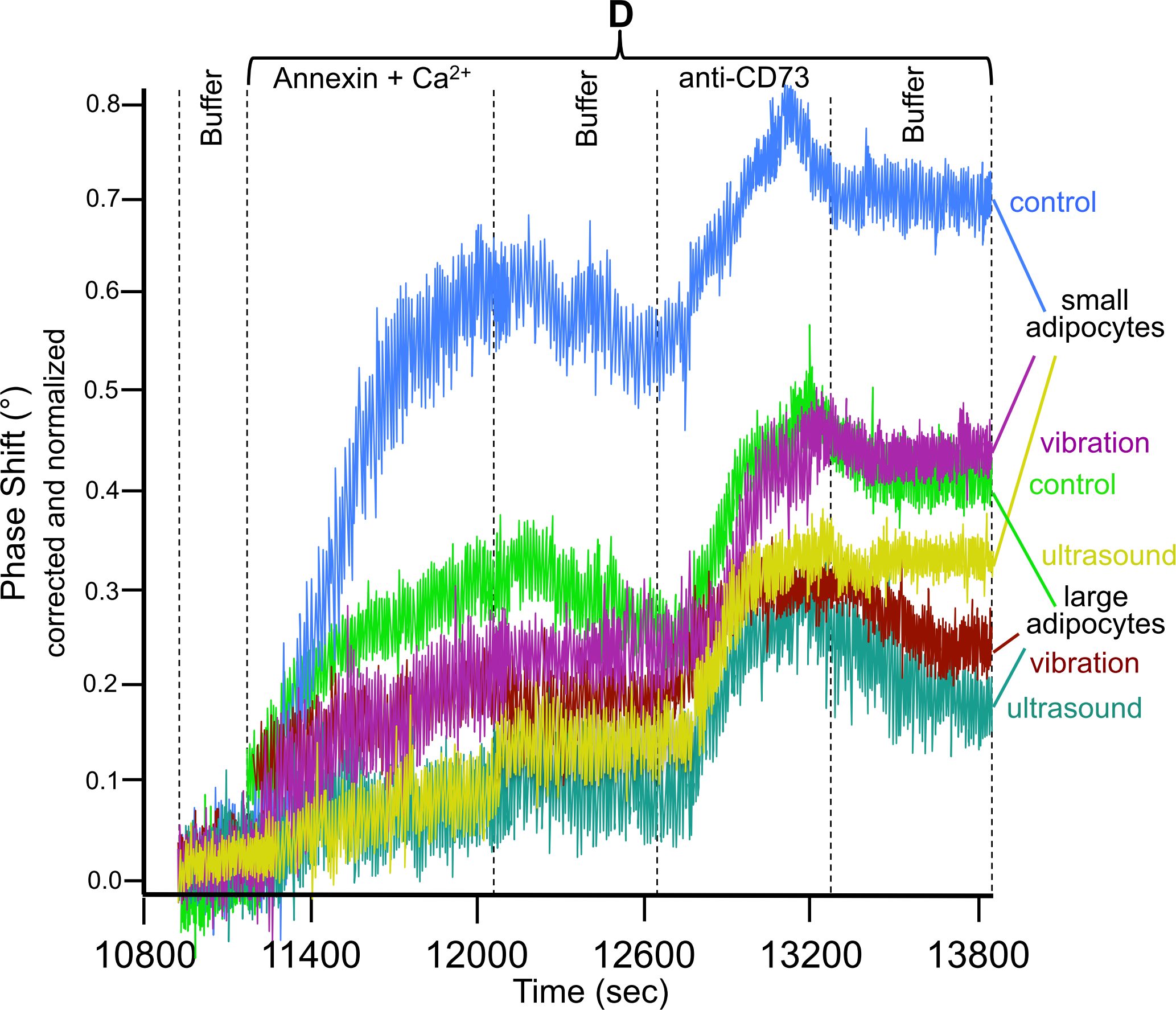
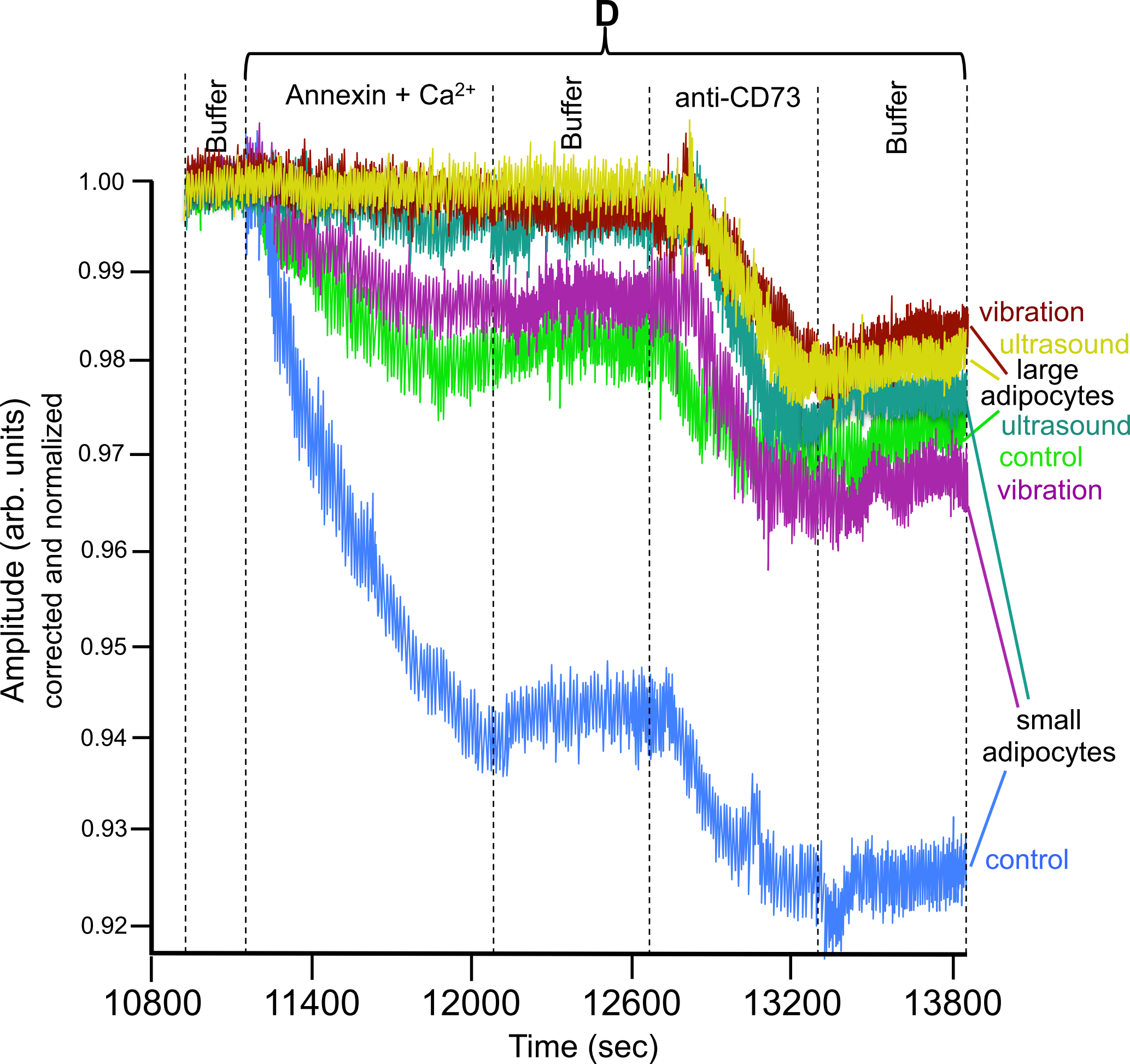
Effect of vibrations and ultrasonic waves on plasma membrane-derived GLEC. The experiment was performed with plasma membranes from small (blue, green, pink curves) and large (yellow, brown, turquoise curves) adipocytes as described for **Figure 2**, however, using a flow rate of 200 μl/min during period B (3000-10200 sec). At the end of period B, the chips were removed from the instrument without emptying of the channels and put into sealed and fitted plastic chambers. Chips with captured GLEC from small or large adipocytes were exposed to ten cycles of vibration (20-sec treatments at max. speed with 20-sec intervals each; neoLab Vortex Mixer 7-2020; pink, brown lines) or three cycles of ultrasound treatment (2-sec bursts at 80 W and 35 kHz with 20-sec intervals at 4°C; Bandelin electronic RF 100; yellow and turquoise lines). Other chips were left untreated as controls (green, blue curves). Phase shift (**a,** as °) and amplitude (**b**, as arbitrary units) were measured during period D and corrected by subtracting the values of the “blank” channel and normalization (set at 0 and 1, respectively, for 10800 sec). Representative overlay diagrams from four independent runs for each condition and adipocyte size (period D shown only) are shown performed with the same chip (and channels re-used four times) for each adipocyte size with two distinct chips each.

**Figure 5.**
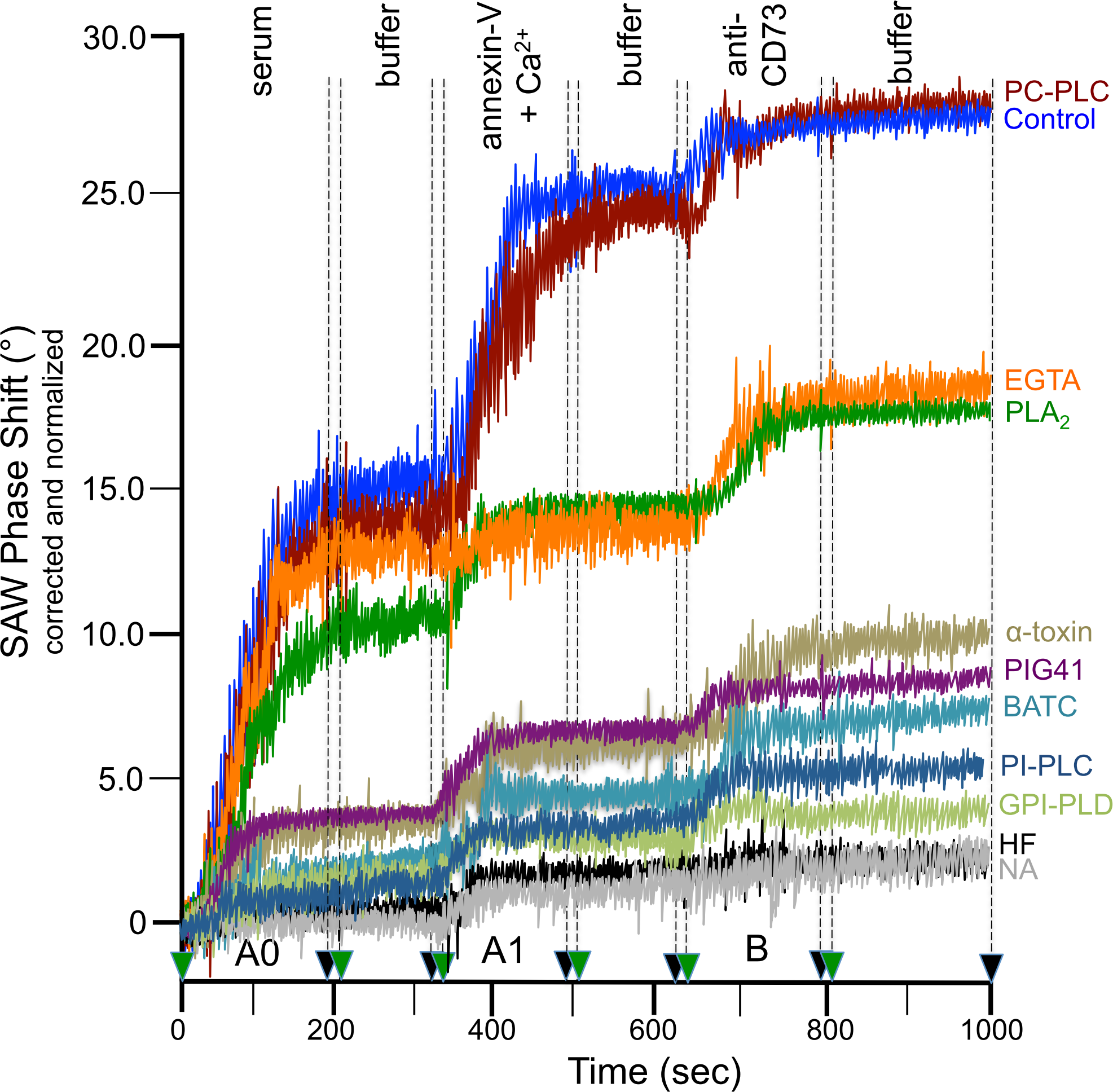
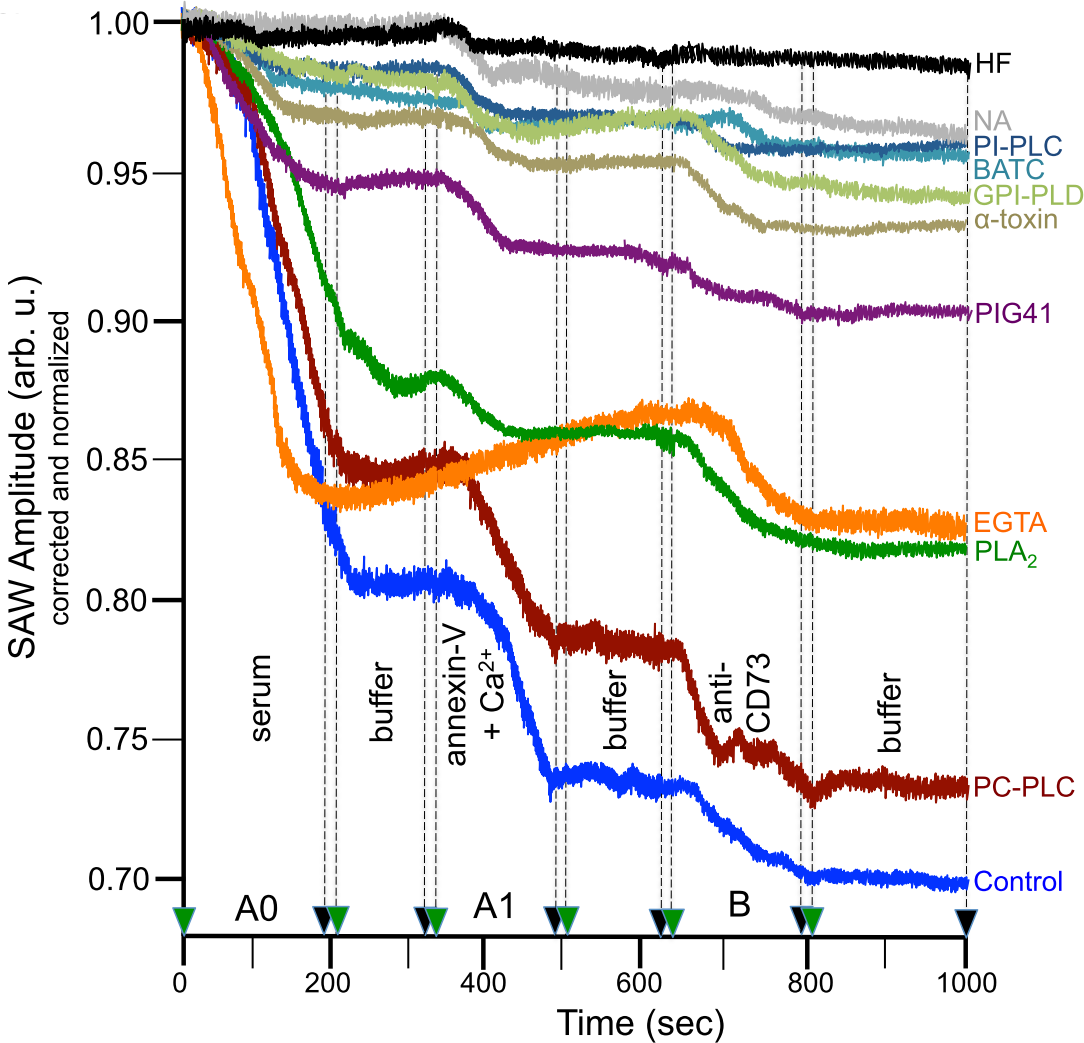
SAW chip-based sensing of GLEC in human serum. 40 µl of pooled serum sample from normal probands (A, B) and diabetic patients (C, D)(**a**) or of serum sample from a diabetic patient (C)(**b**) diluted five-fold with PBS was injected into α-toxin-coated or “blank” channels at a flow rate of 60 µl/min and at 20°C using running buffer (period A0). Prior to the injection, the samples were kept untreated (Control) or incubated (2 h, 30°C) with PC-PLC from *Chlostridium perfringens* (10.0 mU/ml) or with EGTA (2 mM) or with PLA_2_ from honey bee venom (2.5 mU/ml), 0.02% BSA, 144 mM NaCl, 2 mM Ca^2+^, 0.5 mM DTT and 0.2 mM PMSF or with α-toxin-coupled microspheres (subsequent centrifugation; see online methods) or with PIG41 (200 µM) or with BATC (1%) and 144 mM NaCl or with PI-PLC (0.25 mUml) from *Bacillus cereus* or with human GPI-PLD (0.075 mU/ml) or with HF or with NA (see online methods). To demonstrate stable capturing of GLEC by the chip, 240 µl of running buffer were injected subsequently into the channels at a flow rate of 120 µl/min and 37°C. Thereafter, 270 µl of 400 nM annexin-V containing 40 µM Ca^2+^ were injected at a flow rate of 90 µl/min (period A1). To demonstrate stable detection of GLEC by the chip, 240 µl of running buffer were injected subsequently into the channels at a flow rate of 120 µl/min and 37°C. Thereafter, 52 µl of 200 nM anti-CD73 antibodies were injected at a flow rate of 26 µl/min and 37°C (period B). To demonstrate stable detection of GLEC “in sandwich”, 400 µl of running buffer were injected subsequently into the channels at a flow rate of 120 µl/min and 37°C. Phase shift (**a,** as °) and amplitude (**b**, as arbitrary units) are given upon correction for the “blank” and “albumin” channels and normalization (set at 0 for start of the injection at 0 sec after the pretreatments) as original data. Representative overlay diagrams from two independent pretreatments of the same serum samples and three independent measurements using distinct chips are shown.

Next the chips were used in a “lab-on-the-chip” configuration to follow the putative release of GLEC from plasma membranes of releasing cells *in vitro* after their immobilization on the chip gold surface (**Fig. 2a, b**). The stepwise increases in phase shift with each injection of isolated adipocyte plasma membranes, but only in the presence of Ca^2+^ for neutralization of the negative surface charge (period A, blue *vs*. green curves), revealed the apparent hydrophobic interaction of the membranes with the chip. The incremental phase shifts decreased with the number of injections. This is compatible with saturation of a limited number of sites accessible for immobilization. Importantly, only minor detachment of the immobilized plasma membranes (“bleeding”) during the multiple injection and washing (buffer) cycles (period A) was observed as reflected in the only small phase shift decreases. In the following, saturating amounts of plasma membranes from the different preparations with regard to maximal phase shift were used for capturing (period A) which ensured the comparison of roughly identical amounts of immobilized plasma membranes as putative sources for the GLEC. The pronounced phase shift upon injection of anti-insulin receptor-α antibodies into the chip (**Fig. 2a**, end of period A) confirmed the immobilization of adipocyte plasma membranes, which typically express insulin receptor, on the chip surface. The increases in phase shift upon sequential injections of plasma membranes and anti-insulin receptor-α antibodies did not significantly differ between small and large adipocytes (**Fig. 2a, b**, period A) confirming immobilisation of similar amounts of membranes on the chip. Successful immobilization of negatively charged liposomes was reported previously, but interpreted as the formation of flat tethered phospholipid bilayers rather than of (sealed) vesicles on the chip surface^13^. At present, this possibility can not be excluded for the immobilized adipocyte plasma membranes.

For monitoring of the release of GLEC from the immobilized plasma membranes during the subsequent injection of running buffer in the presence of 30 µM PIG37 (period B) they have to be captured by the α-toxin-coated chip simultaneously (period B) and then the plasma membranes detached from the chip by deprotonization (high pH) and chelating of Ca^2+^ (EGTA) for reduction of the phase shift from the plasma membrane-induced to almost basal values (period C). After washing (period C), the stepwise increases in phase shift upon sequential injection of annexin-V and anti-CD73 antibodies, but not anti-insulin receptor-α antibodies, demonstrated the precedent quantitative detachment of the plasma membranes (period B) as well as the subsequent detection of the captured GLEC (period D). Interestingly, the phase shift increases were considerably more pronounced for plasma membranes from small (**a**) compared to large (**b**) adipocytes (period D, blue curves). This differential sensing of GLEC, which may reflect differences in their spec. mass, size and/or amount, was considerably impaired by omission of PIG37 during the simultaneous release and capturing of the GLEC (period B, data not shown).

The specificity of capturing of the plasma membrane-derived GLEC by the chip was confirmed by final injection of 200 µM PIG41 (period D), which caused lowering of the phase shift by roughly 70%. The remaining, apparently unspecific (i.e. not GPI-mediated) phase shift was completely abrogated by excess of mannose (period D). Therefore, only the portion competed for by PIG41 is regarded as GLEC-specific phase shift in the following. Additional evidence for the specificity of chip-based sensing of plasma membrane-derived GLEC was provided by injection of PIG41 (turquoise curves) or mannose (yellow curves) immediately following detachment of the GLEC (period C), but prior to their detection by annexin-V and anti-CD73 antibodies (period D) as well as by use of “blank” chips lacking α-toxin (red curves) since each procedure completely prevented phase shift increase.

Interestingly, injection of the detergent BATC, which is known to preferentially solubilize GPI-AP *vs*. transmembrane proteins from eukaryotic plasma membranes^14^, after detachment of the plasma membranes (**Fig. 2a, b**, period C, black and pink curves) reduced the annexin-V-induced phase shift increases in concentration-dependent fashion, but left unaltered those provoked by anti-CD73 antibodies. This indicates the conversion of GLEC into GPI-AP-BATC micelles. Loss of phospholipids from GLEC was also achieved by bee venom phospholipase PLA_2_ (green curves), capable of cleaving off fatty acids from membrane-associated (glyco)phospholipids including GPI, leaving captured GPI-AP with attached lysophosphatidate moiety only, which were detected by anti-CD73 antibodies, but not by annexin-V. These data are compatible with the proposed composition (GPI-AP, phospholipids) of GLEC. Moreover, plasma membrane-derived GLEC from small (**a**) and large (**b**) adipocytes seem to be structurally related since the shapes of the phase shift curves reflecting detection of the captured GLEC (period D) looked very similar. The more pronounced phase shifts (**Fig. 2a**, **Supplementary Fig. 6a**) and amplitude reductions (**Supplementary Fig. 6b**) for small compared to large adipocytes, which were maintained consistently during each injection (period D), hint to higher spec. mass, size or amount and viscosity, respectively, of GLEC released from plasma membranes of small compared to large cells *in vitro*.

Using the “lab-on-the-chip”, the exogenous and endogenous factors critical for the release of adipocyte plasma membrane-derived GLEC were characterized. Phase shift (**Supplementary Figs. 6a**, **c**) as well as amplitude reduction (**b**, **d**) induced by the GLEC turned out to be positively correlated to the flow rate of the buffer streaming through the microfluidic chip channels for both small and large adipocytes. These findings suggest upregulation of the spec. mass, amount or size and viscosity of the adipocyte plasma membrane-derived GLEC by increase in velocity of the passing extracellular fluid.

Moreover, the adipocyte plasma membranes, while remaining immobilized on the chip, were exposed to several agents known to interfere with membrane integrity (**Supplementary Fig. 7**). Detergent solubilisation (BATC), extraction of fatty acids (BSA) and cholesterol (nystatin) and cleavage of lipidic membrane constituents (PC-PLC, GPI-PLD, PLA_2_), led to altered phase shift (**a**) and amplitude (**b**) in differential fashion compatible with the presence of PC and other phospholipids, GPI and other glycophospholipids, fatty acids as well as cholesterol in adipocyte plasma membrane-derived GLEC. Despite those artificial treatments, it is tempting to speculate that extrinsic “membrane-active” factors play a role in the release and structure of GLEC, a physiologically relevant one possibly represented by serum albumin.

Next it was investigated whether GLEC, similar but not restricted to those released from primary rat adipocytes into the incubation medium and from rat adipocyte plasma membranes in the “lab-on-the-chip” (**Figs. 1**, **2**, see ref. 15 for non-vesicular GLEC subtypes from non-adipose tissues as deduced from indirect experimental evidence), can be detected in rat serum by SAW chip-based sensing. The typical analysis of the serum secretome involves procedures which deleterious effects on the GLEC (e.g. loss as pelleted or floating materials in course of centrifugation, dissociation in course of SDS-PAGE, ELISA or MS) and may suffer from inadequate sensitivity, resolution and throughput (required for clinical studies).

Serum from rats of different genotype and body weight (**Supplementary Table 2**) were used for SAW chip-based sensing of GLEC (**Supplementary Fig. 8a**). Serum from lean Wistar rats provoked considerable and stable (in course of washing) increases in phase shift upon capturing by α-toxin-coated (green curve) *vs*. “blank” (red curve) chips, as well as upon subsequent sequential binding “in sandwich” of annexin-V and anti-CD73 antibodies. However, with this experimental design consistent differential responses for rats differing in both genotype and body weight (e.g. lean Wistar *vs*. obese ZDF rats) were not obtained (**b**). Importantly, a considerably diminished increase in phase shift for obese ZDF compared to lean Wistar rats (**c**) or a more pronounced reduction in amplitude for obese ZF compared to lean Wistar rats (**d**) was achieved by addition of PIG37 at midst of GLEC capturing, which was maintained during subsequent phospholipid detection by annexin-V binding and during washing. The PIG37-dependent difference in amplitude reduction as well as phase shift was found to be maximal with 30 μM as shown for serum from lean ZF *vs*. obese Wistar rats (**e**) and lean ZDF *vs*. obese Wistar rats (**f**), respectively, during consecutive cycles of capturing in the presence of increasing concentrations of PIG37 followed by phospholipid detection and complete displacement of the GLEC from the chip by PIG41.

In order to reduce the number of injection cycles, the capturing and detection steps were combined (**Supplementary Fig. 8g**) which turned out to exert little effect on the serum-induced maximal increase in phase shift and reduction in amplitude (period AB) compared to sequential injections of serum and annexin-V (periods A and B). Consequently, for the chip-based sensing of rat sera simultaneous injection of serum and annexin-V for capturing of the GLEC and detection of the phospholipids (period A1/2) with subsequent measurement of phase shift and amplitude at the end of injection of PIG37 (period B1/2) before regeneration of the chips (periods C-E) was performed (**Fig. 3**). This protocol was most efficient for pairwise differentiation as shown (**a**) for obese ZDF *vs*. obese ZF/Wistar rats (upper panel) and obese ZF *vs*. obese ZDF/Wistar rats (lower panel) with low variance between two consecutive cycles (periods A1-D1 and A2-D2). The differences in phase shift and amplitude between the serum samples in the pairwise combinations were found to be significantly higher and of lower variance when the measurements were performed after the PIG37 injection (**b,** periods B1/2) compared to before (**b**, period A1/2). In conclusion, measurement of both phase shift and amplitude exerted in the presence of PIG37 is required for mutual differentiation of serum from rats of different genotype with similar (obese) body weight (**Figs. 3a**, **b**). In addition, it became apparent that re-use of the chips after sequential displacement of the GLEC using PIG41 (period C1/2), removal of phospholipids using NP-40 (period D1/2) and final washing (period E) is feasible as manifested in non-significant deviations between B1 and B2 (**a**).

After determination of the sensitivity and linearity for chip-based sensing of GLEC with regard to sample volume (**Supplementary Fig. 9a**) and of the variance between (i) the same channel (for re-use), (ii) distinct channels of the same chip and (iii) distinct chips (**b**), appropriate conditions (40 μl sample volume, six re-uses of the same channel) were used for the following measurements of individual and pooled serum samples of eight lean and obese Wistar, ZF and ZDF rats covering different metabolic states (**Supplementary Table 2**) in pairwise comparisons between animals of different genotype which are either lean or obese (**Fig. 3c-h, Supplementary Fig. 10**) and *vice versa* between lean and obese animals which are of identical genotype (**Supplementary Fig. 11a-c**).

In pairwise comparisons of either lean or obese rats, significant differences in phase shift or amplitude were observed for the means of the individually measured serum samples (M) between Wistar and ZDF (**Figs. 3c**, **d**) or ZF (**e**, **f**) rats. Between ZF and ZDF rats trends were monitored only for the obese (**g**, **h**), but not for the lean animals (**Supplementary Fig. 10**). In pairwise comparisons of rats of either Wistar (**Supplementary Fig. 11a**) or ZF (**b**) or ZDF (**c**) genotype trends in phase shift or amplitude were measured for the means of the individual serum samples between lean and obese animals.

In agreement, differences in amplitude were observed between Wistar rats which had been subjected to bariatric or sham surgery and subsequently administered (normal or vitamin-supplemented) high-fat diet^16^ reflecting the acquired lean and obese phenotype, respectively (**Supplementary Table 3**). Consistent differences in amplitude as well as body weight between the type of bariatric surgery (RYGB, VSG) were not recognized. This0020indicates that the body weight loss *per se* was responsible for the surgery-induced increases in amplitude, which in most cases enabled classification of the individual rats as lean or obese using threshold values. Since significant differences in phase shift were measured neither between lean and obese Wistar rats (**Supplementary Fig. 11a**) nor between high-fat diet-fed rats with bariatric and sham surgery (data not shown), the structure of the GLEC, as manifested in altered viscosity, rather than their spec. mass, size or amount is presumably affected by the body weight.

In conclusion, the means of the serum samples (M) as well as the values measured for the pooled samples (P) enabled the differentiation of individual rats for the majority of pairwise comparisons on the basis of phase shifts and amplitudes below or above of the corresponding M/P - 1xSD and M/P + 1xSD, respectively, determined for the counterpart rats (**Supplementary Fig. 12**). Importantly, repetition of this experiment using the same samples, but a distinct instrument (**Supplementary Table 4**) led to similar M and P values which enabled differentiation of the rats according to genotype/body weight with comparable accuracy as was true for the original instrument.

For characterization of the biochemical/-physical nature of the GLEC, serum samples were subjected to (i) lipolytic or chemical (hydrogen fluoride dephosphorylation, nitrous acid deamination) treatments for cleavage of the GPI anchor (**Supplementary Figs. 13**, **14a**, **b**, **c**), (ii) adsorption to microspheres coupled to α-toxin or (c)AMP for depletion of GLEC equipped with GPI-AP or (c)AMP-binding proteins, such as the GPI-AP CD73 and Gce1, respectively (**Supplementary Figs. 14a**, **d**), and (iii) various detergents (**a**, **e**) or cholesterol-extracting agents (**Supplementary Table 5**) for release of lipidic constituents from the GLEC. The observed concentration-dependent declines in both phase shift and amplitude upon treatment with GPI-PLD at moderate concentrations (**Supplementary Fig. 13**) can be explained by partial lipolytic removal of the GPI-AP coat resulting in decreased spec. mass/size and elevated viscosity of the GLEC. At high concentrations, GLEC capturing is prevented in course of complete degradation of the GPI-AP which led to complete restoration of the amplitude. The other treatments resulted in reductions in phase shift which were accompanied by elevations in amplitude (**Supplementary Fig. 14d**) or in elimination of the differences in phase shift and amplitude between serum samples as measured in pairwise combinations (**Supplementary Figs. 14a**, **b**, **c**, **e**). Moreover, incubation of sera with cholesterol-extracting agents prior to measurement provoked significant declines in phase shift and amplitude reduction, which were dependent on the genotype and bodyweight of the rats and the type of agent (**Supplementary Table 5**). This finding is compatible with the presence of cholesterol in serum GLEC and variation of its level with the metabolic state of the rats. Taken together, the data confirm that rat serum contains GLEC which are constituted by GPI-AP, among them CD73 and Gce1, phospholipids and cholesterol and responsible for the differentiation of rats according to the metabolic geno-/phenotype using chip-based sensing.

For an initial characterization of the structure of GLEC, serum samples were exposed to physical stress (**Supplementary Fig. 15**), such as freezing and thawing (**a**), elevated temperature (**b**), centrifugal forces (**c**), ultrasonic waves (**d**) and vibration (**e**). The observed diminuations of the differences in phase shift and amplitude between obese Wistar and ZDF/ZF rats were correlated to the strength of each treatment and thus hint to the operation of weak secondary interactions between the GLEC protein and lipid constituents. Moreover, the pronounced physical instability of rat serum GLEC under conditions which only marginally affected the effects of isolated adipocyte EV or adipocyte incubation medium (containing EV) as manifested in only slightly compromised phase shifts and amplitude reductions (**f**) argues for non-identity of serum GLEC and typical EV^8^.

The apparent physical lability of serum GLEC offered the possibility to correct the total values for phase shift and amplitude for non-GLEC-mediated contributions which may be elicited by captured EV or lipolytically cleaved GPI-AP (**Supplementary Fig. 16**). Calculation of the GLEC-mediated effects as the difference (open bars) between the total (filled bars) and the non-GLEC-mediated alterations (since unaffected by vibration, hatched bars) in phase shift (**a**) and amplitude (**b**) enabled the consistent differentiation of the sera according to genotype and body weight. Importantly, this procedure redundantizes the pairwise comparison of the serum samples (see **Fig. 3**). This protocol together with the limited expenditure for preparation of adequate chips, which relies on (i) their regeneration and multiple use (by competitive displacement of the GLEC from α-toxin rather than its de-/renaturation), (ii) the reproducibility using distinct chips (**Supplementary Fig. 9b**) and instruments (**Supplementary Table 4**) and (iii) the stability and ease of production of the capturing molecule in combination with knowledge about preparation and storage of (serum) samples (**Supplementary Fig. 15**) and compatibility of the SAW sensor technology with analysis of labile macromolecular complexes as reported previously^17^ may enable throughput and routine chip-based sensing of serum GLEC in longitudinal studies to evaluate the possibility that their spec. mass, size, amount and/or viscoelasticity is predictive for the development of metabolic diseases.

Next it was investigated whether the rat serum GLEC measured by chip-based sensing and the adipocyte plasma membrane-derived GLEC identified by the “lab-on-the-chip” are related (**Fig. 4**). In fact, exposure of “lab-on-the-chips” with captured GLEC (during period B) to vibration (pink and brown curves) and ultrasonication (yellow and turquoise curves) after removal from the instrument (period C) considerably reduced the phase shift increases (**a**) and amplitude reductions (**b**) provoked by injection of annexin-V compared to control (blue and green curves). In contrast, the mechanical forces exerted only minor impairments on the binding of anti-CD73 antibodies indicating that the capturing of GPI-AP *per se* remained unaffected. These findings argue for structural similarity between rat adipocyte plasma membrane-derived GLEC and rat serum GLEC with both representing labile complexes of GPI-AP, phospholipids and cholesterol.

In addition to mechanical forces (e.g. buffer flow) and “membrane-active” agents (e.g. detergents, enzymes, albumin in the buffer), the metabolic state of the rats as donors for the adipocyte plasma membrane-derived GLEC may critically determine their release in the “lab-on-the-chip”. In fact, both specific phase shift (**Supplementary Fig. 17a**) and amplitude reduction (**b**) were significantly reduced for total adipocytes from obese *vs*. lean ZDF rat and old *vs*. young Wistar rats as well as for small adipocytes from young *vs*. large adipocytes from old Wistar rats. Remarkably, isolated adipocyte plasma membranes of obese and old Wistar rats were differentiated from one another on basis of the specific phase shift induced by the GLEC upon release from them (**a**). Thus adipocyte plasma membrane-derived and serum GLEC, as monitored by chip-based sensing, are affected by the metabolic state of the corresponding donor rats in comparable fashion.

The presence of BSA during release of the plasma membrane-derived GLEC (**Supplementary Fig. 18**) led to significant increases in the differences in specific phase shift (**a**) and amplitude reduction (**b**) for total adipocytes from between obese *vs*. lean ZDF rats as well as for small adipocytes from young *vs*. large adipocytes from old Wistar rats. On basis of the data obtained with the chip-based sensing *in vitro* (isolated plasma membranes and primary adipocytes) and *in vivo* (serum) it is tempting to speculate that *in vivo* (i) release of the GLEC with regard to spec. mass, size, amount and viscoelasticity is determined by the extracellular fluid (blood pressure, albumin, lipases) and the metabolic state of the releasing cells and (ii) plasma membranes of adipocytes can operate as source for serum GLEC. However, the presence and abundance of adipocyte-derived GLEC in serum remain to be investigated.

Albeit the expression of GLEC *in vivo* has not been reported so far, alterations in the specific as well as total expression pattern of GPI-AP and phospholipids in plasma, as measured by ELISA (for CD59), proteomics and lipidomics, have been described for aging^18^, breast cancer^19^ and Alzheimer’s disease^20^, respectively. However, changes in the GPI-AP-to-cholesterol-to-phospholipid ratio of (serum and plasma membrane-derived) GLEC with the corresponding insults on spec. mass, size, amount and viscoelasticity, which apparently differ between normal and hyperinsulinemic/ hyperglycemic rats (**Supplementary Table 6**) escape detection by “single-parameter” methods, such as MS. Moreover, GLEC are probably heterogenous in structure depending on their cellular origin and the (patho)physiological mechanisms of their release and degradation, which remain a matter of speculation at present (**Supplementary Fig. 19**), in particular the putative link to the normoinsulinemic/glycemic (**a**) compared to the hyperinsulinemic/glycemic state (**b**).

Finally, the possibility of expression of GLEC in human serum similar to those described above for rat serum was tested by SAW chip-based sensing (**Fig. 5**). Sequential injection of a pooled (**a**) or individual (**b**) human serum sample (period A0), annexin-V in the presence of Ca^2+^ (period A1) and anti-CD73 antibodies (period B) into α-toxin-coated channels elicited considerable increases in phase shift (**a**) and reductions in amplitude (**b**), which each resisted subsequent washing (Control). In contrast, only very minor effects on phase shift and amplitude were observed with “blank” channels (data not shown). The annexin-V/Ca^2+^-, but hardly the serum- and anti-CD73-induced upregulations of phase shift and amplitude reduction were abrogated by EGTA and PLA_2_ (but not by PC-PLC), demonstrating the specificity of GLEC-associated phosphatidylserine detection by annexin-V. The specificity of capturing of the GLEC was confirmed by drastically diminished upregulations of phase shift (**a**) and amplitude reduction (**b**) compared to control with serum which had been depleted of GPI-AP by adsorption to α-toxin-coupled microspheres prior to injection or the inclusion of PIG41 during the injection. This was even more pronounced with pretreatment of the samples with BATC prior to injection which is compatible with the presence of phospholipids, GPI lipids and GPI-AP in the GLEC and their solubilization by BATC. The presence of GPI lipids and GPI-AP in the GLEC was confirmed by reductions in the phase shift increase (**a**) and amplitude reduction (**b**) by 80 to 90% in course of enzymic and chemical pretreatments with PI-PLC or GPI-PLD and HF or NA, respectively, which all are known to specifically cleave within the phospholipid or core glycan portions of GPI-AP.

Taken together, these data strongly argue for the expression of complexes consisting of GPI-AP and phospholipids in human serum which closely resemble the rat serum and adipocyte plasma membrane-derived GLEC. However, in contrast to the observed correlation between the genotype/body weight and the PIG37-dependent alterations in phase shift and amplitude during chip-based sensing of rat serum GLEC, the analysis of fresh unfrozen serum from ten probands did not reveal clear-cut similarities in the kinetics of phase shift and amplitude changes, i.e. shapes or patterns of the corresponding curves, for the period of capturing as well as detection of the GLEC (**Supplementary Fig. 20**), even in the presence of PIG37 during capturing for maximal differentiation (**Supplementary Table** 7), for both control subjects and diabetic patients. Moreover, no consistent differences between them could be identified. Remarkably, human serum GLEC exhibited higher stability towards physical stress compared to rat GLEC (**Supplementary Figs. 21, 22**), which prevented the determination of threshold values for specific GLEC- induced phase shift and amplitude as is feasible for rats (**Supplementary Fig. 16**). Thus it can not be excluded that in human serum subpopulations of GLEC are expressed which correlate with the metabolic state, but are of lower stability compared to rat GLEC (and the human GLEC described above) and become lost during the currently used procedures of sample processing and sensing. Alternatively, the number of samples analyzed could have been inadequate to elucidate patterns for chip-based sensing of serum GLEC for metabolically differing probands. The possibility that GLEC in human serum are indicative for different metabolic (e.g. diabetic) states deserves further investigation.

## METHODS

Methods, including statements of data and material availability and any associated references are available in the online version of the paper.

## ACKNOWLEDGEMENTS

This work was supported in part by funding to M.H.T. from the Alexander von Humboldt Foundation, the Helmholtz Alliance *ICEMED* & the Helmholtz Initiative on Personalized Medicine *iMed* by Helmholtz Association, and Helmholtz cross-program topic “Metabolic Dysfunction”. We thank M. Bauer and P. Kotzbeck (Helmholtz Zentrum Munich, Institute for Diabetes and Obesity) for helpful discussions regarding EV. We also appreciate the collaborative help and experimental advice of Thomas Gronewold (SAW Inc.) for implementation of the SAW sensor technology.

## AUTHOR CONTRIBUTIONS

G.A.M. conceived the project and designed and performed the experiments; A.W.H., K.S. and A.L. provided the serum samples; G.A.M., K.S. and A.W.H. analysed the data; G.A.M., A.W.H., K.S., A.L. and M.H.T. discussed the data; G.A.M. wrote the paper with input from all coauthors; M.H.T. supervised the work.

## ONLINE METHODS

### Materials

ß-amidotaurocholate (BATC) was synthesized according to published methods^21^. NBD-FA was from Molecular Probes Inc. (Eugene, OR, USA). Collagenase (Worthington, CLS type I, 190-250 units/mg) was provided by Biochrom (Berlin, Germany). Human recombinant insulin was supplied by the biotechnology department of Sanofi Pharma Germany (Frankfurt am Main, Germany). Partially purified phosphatidylinositol-specific phospholipase C (PI-PLC) from *Bacillus cereus* (1 unit/mg protein), phosphatidylcholine-specific phospholipase C (PC-PLC) from *Clostridium perfringens* (25 units/mg protein), phospholipase A_2_ (PLA_2_) from honey bee venom (*Apis mellifera*)(1800 units/mg), bovine serum albumin (BSA; fraction V, defatted), rat serum albumin (RSA), phenylisopropyladensine (PIA), adenosine deaminase (ADA), PMSF, purified α- mannosidase from *Canavalla ensiformis* (proteomics grade), saponin, digitonin, filipin, nystatin and cAMP-agarose were obtained from Sigma-Aldrich Chemie GmbH (Munich, Germany). Recombinant (*E. coli*) human glycosylphosphatidylinositol-specific phospholipase D1 (GPI-PLD1)(0.5 units/mg protein) was purchased from Creative BioMart Inc. (New York, USA). Streptolysin-O was bought from VWR Scientific. 5’-AMP-agarose was delivered by Jena Bioscience GmbH (Jena, Germany). Anti-CD73 antibodies (rabbit polyclonal, affinity-purified, IgG isotype, prepared against recombinant full-size human CD73 with reactivity against rat CD73) were obtained from Genetex/Biozol (Eching, Germany). Annexin-V (human, recombinant) was purchased from ProSpec (East Brunswick, NJ, USA). Recombinant (*E. coli*) *N*-glycanase F from *Flavobacterium meningosepticum*, recombinant (*Pichia pastoris*) proteinase K (PCR grade), protease inhibitor cocktail (“complete ULTRA” tablets), octylglucoside (OG) and Nonidet-P40 (NP-40) were provided by Roche Biochemicals (Mannheim, Germany). GF/C glass fiber filters were delivered by Whatman (Maidstone, UK). 5’-AMP-Sepharose beads were purchased from LKB/Pharmacia (Freiburg, Germany). “Stains-all” was provided by ICN Biochemicals (Cleveland, OH, USA). 1-ethyl-3-[3-dimethylaminopropyl]carbodiimide (EDC) and N-hydroxysulfosuccinimide (Sulfo-NHS, premium grade) were bought from Pierce/Thermo Scientific (Rockford, IL, USA). Tetrahydrofurane (THF), PBS (tablets), TRIS, HEPES, MES, silver nitrate, thin layer chromatography (TLC) plates (Si-60), dinonylphtalate, ethanolamine, mannose, Tween, glycine and all other reagents (highest purity available) were from Merck (Darmstadt, Germany).

### Chemical synthesis of phosphoinositolglycans (PIG) and (G)PI-PLC inhibitor GPI- 2350

The methods for the synthesis of PIG41 and PIG37 have been described previously^12^. In brief, the glycosidic linkages have been made stereoselective by the trichloroacetimidiate methodology. For introduction of the phosphate, the tetrabenzylpyrophosphate/sodium hydride or the phosphitylation/oxidation protocol was used. For introduction of the sulfate, a solution of trimethylamine-sulfur trioxide complex in pyridine was used. The structure of the PIG was characterized by mass, ^1^H NMR und ^31^P NMR spectroscopy. The method for the synthesis of GPI-2350 has been outlined in detail previously^22^. In brief, racemic 1,4,5,6-tetra-*O*-benzyl-*myo*-inositol was phosphorylated with dodecyl-phosphonic dichloride in the presence of triethylamine and dimethylaminopyridine. The resulting tetra-*O*-benzyl-*myo*-inositol-1,2-cyclo-dodecyl-phosphonic acid was debenzylated by catalytic hydrogenation on charcoal to yield *myo*-inositol-1,2- cyclo-dodecyl-phosphonic acid (GPI-2350) as a mixture of diastereomeres.

### Preparation of α-toxin

α-Toxin was purified from the culture supernatant of *Clostridium septicum* (strain KZ1003) after an 18-h culture in brain-heart infusion broth (Difco). After precipitation by 60% saturated ammonium sulfate and centrifugation, the pellet was dissolved in 10 mM sodium phosphate (pH 7.0) and then subjected to cation exchange chromatography on SP-Toyopearll 650M^23^. After elution, the corresponding fraction was again precipitated by 60% saturated ammonium sulfate and centrifuged. SDS-PAGE and Coomassie-staining of the pellet materials resulted in a single protein band at a position corresponding to a MW of 48 kDa. α-Toxin was suspended in 100 mM MES/KOH (pH 6.5) at 1 mg/ml.

### Animal handling

16 male Wistar rats (Crl:WI(WU)), 16 male Zucker diabetic fatty rats (ZDF-*Lepr*^*fa*^/Crl) and 15 male Zucker fatty rats (Crl:ZUC(Orl)-*Lepr*^*fa*^/Zucker)^24^, respectively, were obtained from Charles River (Sulzfeld, Germany). The male ZDF rats were 14 weeks old at start of the induction of the lean and obese phenotype, respectively. At this age, male obese ZDF rats already exhibit a full-blown diabetic phenotype with a catabolic metabolic state. This explains the same body weight range of lean and obese ZDF rats at this age. Rats were housed two *per* cage in an environmentally controlled room with a 12:12-h light–dark circle (light on at 06:00) and *ad libitum* access to food and water. Standard rat chow (17.7 kJ/g, Ssniff diet R/M-H, V1535 with 18% crude protein, 4.7% sugar, and 3.5% crude fat) and a cafeteria diet (21.6 kJ/g, Ssniff diet EF R, 10 mm with 13.5% crude protein, 42.8% sugar, and 22.5% crude fat)(Ssniff, Soest, Germany) were used for the induction of the lean and obese phenotype, respectively, as described previously^25^. All experimental procedures were conducted in accordance with the German Animal Protection Law (paragraph 6) and corresponded to international animal welfare legislation and rules.

### Serum collection

Blood was collected from the tail vein of conscious rats under terminal isoflurane, placed in serum gel tubes and centrifuged (1,000x*g*, 5 min, 4°C) to yield serum, which was immediately frozen in liquid N_2_ and then stored at −20°C. The samples were thawed by incubation at 37°C and then immediately used for measurement unless indicated otherwise. Blood metabolic parameters were determined enzymatically according to standard procedures using commercially available kits (for glucose Glucoquant glucose/HK kit; Roche, Mannheim, Germany) and the Hitachi 912 instrument. Insulin concentration was assayed for 15 µl of sample volume with an enzyme immunoassay (ultrasensitive rat insulin ELISA; Mercodia 10-1248-10, Uppsala, Sweden) covering a sensitivity range of 0.5-100 µg/l and displaying the following crossreactivities: Human insulin 167%, porcine insulin 476%, sheep insulin 179%, bovine insulin 78%. All assays were performed according to the instructions of the manufacturers.

Human serum was collected using standard clinical procedures. Glucose was measured with the aid of the Glucose HK Gen.3 kit (Roche Diagnostics, Mannheim, Germany). Written informed consent was obtained from all study participants, and the protocol was approved by the ethical review committee of the Ludwig-Maximilians-University (**study ID ???**).

### Preparation of isolated rat adipocytes

Isolated rat adipocytes were prepared as described previously^26^ with the following modifications: Rats were killed by cervical dislocation. Epididymal fat pads from male Wistar rats (weight see below, fed ad libitum) were washed several times in Krebs-Ringer-Henseleit buffer (KRH; 25 mM HEPES free acid, 25 mM HEPES sodium salt, 1.2 mM KH_2_PO_4_, 140 mM NaCl, 4.7 mM KCl, 1.2 mM MgSO_4_, 2.5 mM CaCl_2_) supplemented with 0.75% (w/v) BSA and adjusted to pH 7.2 (=KRHLB) and then cut into two to three pieces. Two pieces each were incubated with 1.5 ml of digestion buffer (10 mg collagenase and 9 mg glucose in 10 ml of KRH containing 5% BSA [=KRHHB], 1 mM sodium pyruvate, 5.5 mM glucose, 0.5 U/ml ADA, 200 nM PIA, 100 µg/ml gentamycin, 50 units/ml penicillin and 50 µg/ml streptomycin sulfate) for 15–20 min at 37°C in a shaking water bath (240 cycles/min) at constant bubbling with 95% O_2_/5% CO_2_. Released adipocytes were passed through a nylon web (mesh size 750 *μ*M) with no pressure other than gentle teasing with a plastic spatula for passing of the cells through and then centrifuged (500x*g*, 1 min, swing-out rotor). The supernatant layer of cells was washed three times with 25 ml each of KRH by centrifugation. After final aspiration of the infranatant, the adipocytes were suspended in KRH containing 2% BSA at a lipocrit of 10% corresponding to 100 µl of packed cell volume *per* ml incubation volume (determined by aspiration of small aliquots into capillary hematocrit tubes and centrifugation for 60 sec in a microhematocrit centrifuge in order to determine the fractional occupation of the suspension by the adipocytes). 10% lipocrit corresponds to about 6.5, 1.5 and 0.3×10^6^ adipocytes of small, medium and large size, respectively, *per* ml.

### Size separation of adipocytes

Adipocytes were separated according to size using published procedures^27^ with the following modifications: Adipocytes prepared from 1- month (120-140 g), 2-month (200-240 g) and 6-month (380-420 g) old male rats were collected by flotation (200x*g*, 2 min, 30°C) and then filtered through serial nylon mesh screens with pore sizes of 400, 150 and 75 µm in this order to obtain small adipocytes (diameter < 75 µm from 1-month old rats) from the flow-throughs or 75, 150 and 400 µm in this order to obtain large adipocytes (diameter > 400 µm) from the filter cakes or 400, 75 and 150 µm in this order to obtain middle-sized adipocytes (75 µm < diameter < 150 µm from 2-months old rats) from the flow-through, filter cake and flow-through, respectively. After fixation of aliquots of the adipocyte suspension with osmic acid, cell number was determined using a Coulter counter. The adipocyte suspension was adjusted to the desired titer with adipocyte buffer (20 mM HEPES/KOH, pH 7.4, 140 mM NaCl, 4.7 mM KCl, 2.5 mM CaCl_2_, 1.2 mM MgSO_4_ and 1.2 mM KH_2_PO_4_,) supplemented with 0.5% BSA, 100 µg/ml gentamycin, 50 units/ml penicillin and 50 µg/ml streptomycin sulfate. Total lipid content of the adipocyte suspension (lipocrit) was measured as described above.

### Measurement of insulin sensitivity and responsiveness of adipocytes as insulin- stimulated lipogenesis

Adipocytes were assayed for lipogenesis as described previously^26^ with the following modifications: 200 μl of adipocyte suspension were supplemented with 190 μl of KRHLB containing 1 mM glucose and 10 μl of an insulin solution to yield the desired final concentration and incubated (15 min, 37°C) in a slowly shaking water bath. The assay was started by addition of 50 μl (unless otherwise indicated) of 0.9 mM NBD-FA (prepared daily from a 100-mM stock solution in ethanol by dilution with KRHLB under mild heating). After incubation (90 min, 37°C) under mild shaking (stage 11, thermomixer, Eppendorff, Germany), lipogenesis was terminated by filtration of the total mixtures over GF/C-filters under vacuum. The filters were rapidly washed three times with 1 ml of KRHLB each, placed in 20-ml plastic scintillation vials and extracted with 400 μl THF for 15 min under rigorous shaking. 300 μl of the extract was transferred into new tubes and centrifuged (15,000x*g*, 5 min). The supernatant was dried (SpeedVac) and suspended in 50 μl THF. 5-μl samples were analysed by TLC on silica gel Si-60 plates using 78 ml diethyl ether, 22 ml petrol ether and 1 ml acetic acid as solvent system. The amount of fluorescent lipid products on the dried plates was determined by fluorescence imaging with excitation at 460 nm and emission at 540–560 nm using a phosphorimaging system. The relative peak area of each lipid product were corrected for a background value (derived from an equal-sized region of the TLC plate which does not contain any lipid product).

### Preparation of plasma membranes from rat adipocytes

Plasma membranes were prepared from primary rat adipocytes as described previously^27^ with the following modifications: Adipocytes (10^7^ cells) prepared from epididymal adipose tissue depots were washed and immediately homogenized in 2 ml of lysis buffer (25 mM TRIS/HCl, pH 7.4, 0.5 mM EDTA, 0.25 mM EGTA and 0.25 M sucrose, supplemented with 10 g/ml leupeptin, 2 μM pepstatin, 10 g/ml aprotinin, 5 μM antipain and 200 μM PMSF) using a motor-driven Teflon-in-glass homogenizer (10 strokes with a loosely fitting pestle) at 22°C. The defatted postnuclear infranatant (1,500x*g*, 5 min) was centrifuged (12,000x*g*, 15 min). The pellet was suspended in 1 ml of lysis buffer and layered on top of an 8-ml cushion of 28% Percoll, 0.25 M sucrose, 25 mM TRIS/HCl (pH 7.0) and 1 mM EDTA. After centrifugation (45,000x*g*, 60 min), the purified plasma membranes were withdrawn from the lower third of the gradient (0.5 ml), then pelleted (200,000x*g*, 90 min) and finally suspended in 10 mM HEPES/KOH (pH 7.5), 150 mM NaCl and 100 mM sucrose at 50 μg protein/ml.

### Preparation of incubation medium from rat adipocytes

Adipocytes were suspended in 10 ml of adipocyte buffer (20 mM HEPES/KOH, pH 7.4, 140 mM NaCl, 4.7 mM KCl, 2.5 mM CaCl_2_, 1.2 mM MgSO_4_, 1.2 mM KH_2_PO_4_, 2% (w/v) BSA, 100 μg/ml gentamycin, 1 mM sodium pyruvate and 5.5 mM glucose) at a lipocrit of 0.2% and then incubated (20 h, 37°) in a shaking water bath (100 cycles/min) under constant bubbling with 5% CO_2_/95% O_2_. Thereafter, twelve 350-µl portions of the total mixtures were transferred into microfuge tubes (Beckman) pre-filled with 100 μl of dinonylphtalate and then centrifuged (1,000x*g*, 1 min, 20°C). The tubes were cut through the dinonylphtalate layer separating the adipocytes at the top from the incubation medium in the bottom part of the tubes which was rescued. Care was taken to minimize the volume of dinonylphtalate taken along with the incubation medium in the bottom part. After transfer of the incubation medium (300 μl) into 1-ml Eppendorf cups and washing the bottom part of the tube once with 300 μl of adipocyte buffer containing 0.5 mM DTT and protease inhibitor mix, the combined medium and washing fluid were centrifuged (1,000x*g*, 5 min, 4°C) for the removal of cell debris.

### Preparation of total and Gce1-/CD73-harboring EV from rat adipocytes

EV from isolated rat adipocytes were obtained using published procedures^27^ with the following modifications: For preparation of total EV, three 180-µl portions of the supernatant obtained were centrifuged (Beckman Airfuge, A-110 fixed angle rotor, 30 psig, 110,000 rpm, 30 min, 4°C). After careful aspiration of the supernatants, the pellets were resuspended in 100 μl of EV buffer (10 mM TRIS/HCl, pH 7.4, 250 mM sucrose, 1 mM EDTA, 0.5 mM EGTA, 140 mM NaCl, 1 mM DTT and protease inhibitor cocktail) each (vortexing), combined and re-centrifuged as above. The pellet was washed two times with 100 μl each of the same buffer. After purification by sucrose density gradient centrifugation (200,000x*g*, 18 h, Beckman SW41) on 10-70% sucrose in PBS prepared with Gradient Mate device (BioComp, Fredericton, New Brunswick, Canada), fractions with densities of 1.12-1.22 mg/ml were combined, diluted ten-fold with PBS and centrifuged (100,000x*g*, 2 h, 4°C). The pelleted total EV were suspended in the initial volume of EV buffer and re-centrifuged.

For affinity purification of Gce1- and CD73-harbouring EV, total EV were adsorbed to cAMP-agarose and 5’-AMP-Sepharose, respectively. For this, 100-µl suspensions were supplemented with 200 µl of cAMP-agarose/5’-AMP-Sepharose beads (50 mg each in 1 ml of 100 mM HEPES/KOH, pH 7.4, 140 mM NaCl, 1 mM MgCl_2_, 0.5 mM DTT and protease inhibitor cocktail) and then incubated under continuous head-over rotation (60 cycles *per* min) of the tubes. Thereafter, the incubation mixtures were centrifuged under conditions (1,000x*g*, 5 min, 4°C), which did not lead to sedimentation of unadsorbed EV (lacking Gce1 and CD73) *per se*, but were sufficient for the quantitative sedimentation of the Sepharose beads (with the Gce1-/CD73-harbouring EV adsorbed). The collected bead-EV complexes were washed twice by suspending in 1 ml of EV buffer, each, and centrifugation (1,000x*g*, 5 min, 4°C). The final pellet was dissociated in 100 µl of EV buffer containing 100 µM AMP and cAMP each. After incubation (30 min, 4°C) and centrifugation (12,000x*g*, 5 min, 4°C), the supernatant containing the Gce1-/CD73-harbouring EV was recovered, frozen in liquid N_2_ and stored at −80°C until use.

#### Coupling of α-toxin to microspheres

5.0×10^6^ of the uncoupled magnetic carboxylated microspheres (MagPlex™-C, Luminex Corp.) in a microcentrifuge tube were resuspended according to the instructions of the product information sheet, placed into a magnetic separator and then subjected to separation for 30 to 60 sec. After removal of the supernatant and subsequently of the tube from the separator, the microspheres were resuspended in 100 μl of H_2_O_bidest_ by vortexing and sonication for about 20 sec. Thereafter the tube was again placed into the magnetic separator and separation was allowed to occur for 30 to 60 sec. After removal of the supernatant and subsequently of the tube from the separator, the washed microspheres were resuspended in 80 μl of 100 mM sodium phosphate (pH 6.2) by vortexing and sonication for about 20 sec. After addition of 10 μl of 50 mg/ml Sulfo-NHS (diluted in H_2_O_bidest_) to the microspheres and gentle mixing by vortexing, they were supplemented with 10 μl of 50 mg/ml EDC (diluted in H_2_O_bidest_). The mixture was incubated (20 min, 22°C) under gentle mixing by vortexing at 5-min intervals. Subsequently the tube was placed into the magnetic separator and separation allowed to occur for 30 to 60 sec. The supernatant was removed and then the tube from the separator. The activated microspheres were resuspended in 250 μl of 50 mM MES/KOH (pH 5.0) by vortexing and sonication for about 20 sec. The washing step with magnetic separation and resuspension in 100 mM MES/KOH (pH 6.5) was repeated three times. After the last separation, the microspheres were suspended in 100 μl of 100 mM MES/KOH (pH 6.5) by vortexing and sonication for about 20 sec. 200 μg α-toxin was added to the activated and washed microspheres and the total reaction volume adjusted to 500 μl of 100 mM MES/KOH (pH 6.5). After vortexing, the mixture was incubated (2 h, 22°C) under head-over rotation. Subsequently the tube was placed into the magnetic separator and separation allowed to occur for 30 to 60 sec. The supernatant was removed and then the tube from the separator. The coupled microspheres were resuspended in 500 μl of PBS/TBN (PBS, pH 7.4, 0.1% BSA, 0.02% Tween-20 and 0.05% azide) by vortexing and sonication for 20 sec. After incubation (30 min, 22°C) under head-over rotation, the microspheres were subjected to magnetic separation for 30 to 60 sec. The supernatant was removed and then the tube from the separator. The microspheres were resuspended in 1 ml of PBS/TBN by vortexing and sonication for 20 sec. The washing step with magnetic separation and resuspension was repeated three times with 1 ml each of PBS. After the last separation, coupled and washed microspheres were suspended in 500 μl of PBS/TBN by vortexing and sonication for about 20 sec and then stored at 4°C in the dark.

### Depletion of samples from GPI-harboring entities

#### Medium samples

10 ml of incubation medium were added to 500 μl of PBS containing the microspheres coupled to α-toxin in a 15-ml vial. After vortexing, the mixtures were incubated (30 min, 22°C) under head-over rotation. Subsequently the tubes were placed into the magnetic separator and separation allowed to occur for 30 to 60 sec. The supernatants were removed and then the tube from the separator. The coupled microspheres were resuspended in 500 μl of PBS/TBN by vortexing and sonication for 20 sec. After incubation (30 min, 22°C) under head-over rotation, the microspheres were subjected to magnetic separation for 30 to 60 sec. The supernatant was removed and then the tube from the separator. The microspheres were resuspended in 1 ml of PBS/TBN by vortexing and sonication for 20 sec. The washing step with magnetic separation and resuspension was repeated three times with 1 ml of PBS each. <u>Serum samples</u>: Microspheres (1×10^5^) coupled to α-toxin and resuspended in 10 μl of PBS/TBN were added to 90 μl serum and after vortexing and sonication for 20 sec incubated (30 min, 22°C) under head-over rotation. The microspheres were subjected to magnetic separation for 30 to 60 sec. The supernatant was transferred to a new tube and then the tube removed from the separator. The microspheres were resuspended in 100 μl of PBS/TBN by vortexing and sonication for 20 sec. After the magnetic separation, the supernatant was removed and combined with the initial supernatant.

### Depletion of samples from Gce1 and CD73

cAMP-agarose and 5’-AMP-agarose ([c]AMP-agarose) or agarose alone were suspended in PBS/TBN at 50 mg/ml. 10 μl were added to 90 μl of serum and then incubated (30 min, 22°C) under head-over rotation. Subsequently, the suspensions were subjected to centrifugation (300x*g*, 5 min, 22°C). The supernatants were transferred to new tubes. The beads in the pellets were resuspended in 100 μl of PBS/TBN by gentle vortexing and then re-centrifuged. The supernatants were removed and combined with the initial supernatants.

### Coupling of α-toxin to the chip surface

Coupling reactions were performed as described previously^28-32^ with the following modifications: For the generation of GLEC- capturing chips, α-toxin (in 20 mM TRIS/HCl, pH 7.5, 150 mM NaCl and 10% glycerol) diluted in immobilization buffer (10 mM sodium acetate, pH 5.5) was coupled to the channels of activated long-chain 3D carboxymethyl (CM) dextran chips (SAW Instruments Inc., Bonn, Germany) in a SamX instrument (SAW Instruments Inc.). For this, the surface of microfluidic channels of sensor chips was primed by three injections of 150 μl, each, of immobilization buffer at a flow rate of 45 μl/min. Then the chip surface was activated by a 150-μl injection of 0.2 M EDC and 0.05 M Sulfo-NHS (mixed from 2x-stock solutions right before injection) at a flow rate of 45 μl/min. After a waiting period of 1 min, the coat protein (e.g. α-toxin) was coupled by injection of 200-μl portions at a flow rate of 60 μl/min and subsequent waiting for 2 min. After additional washing with three 150-μl portions of running buffer PBST (PBS containing 0.005% Tween 20 [v/v]) at a flow rate of 30 μl/min and waiting for 5 min, the residual activated groups on the chip surface were capped by injecting 150 μl of 1 M ethanolamine (pH 8.5) at a flow rate of 45 μl/min. For the generation of a “blank” channel lacking coat protein, one channel was activated and blocked with injection of buffer instead of a coat protein. Measurements were performed at 22°C. The flow rates are given in the figure legends or can be derived from the figures considering the start and termination points of the solution flow for sample injections or washing cycles as indicated with green and black arrows, respectively. Chips were regenerated by successive injections of 60 μl of 10 mM glycine (pH 3.5) and 30 μl of 4 M urea with waiting for 5 min after each injection and final injection of 300 μl of regeneration buffer (PBS, pH 7.4, 1 M NaCl, 0.03% Tween and 0.5% glycerol) and 300 μl of PBST and used up to six times without significant loss of capturing (through α-toxin) capacity.

### SAW measurement, instrumentation and evaluation

The Seismos NT.X Instrument for Surface Acoustic Wave (SAW) chip-based sensor (NanoTemper Technologies, Munich, Germany), formerly SamX (SAW Biosensor GmbH, Bonn, Germany) integrates a high-frequency unit, control and reader units and all fluid handling components required for a systematic buffer and analyte solution handling (S-sens K5). This enables fluidic and electrical contacting of the chip with four independent flow-through microfluidic channels at a stable temperature of 22°C (Δ*T* = 0.05 °C by means of four peltier elements). Mass loading and loss of elasticity (gain of viscosity) resulting from biomolecular interaction processes on the chip surface will result in phase shift and amplitude reduction, respectively, of the SAW generated by the inverse piezoelectric effect. The instrument was run and the signals generated were recorded in real-time using a double-frequency measurement mode as described previously^33-38^ with the following modifications: Measurements were performed with a continuous buffer stream at the flow rates and temperatures as indicated. For each chip, the phase shift and amplitude generated by the α-toxin-coated channels were corrected for unspecific non-GPI-mediated interactions by subtraction of the values of the “blank” channel. In addition, in case of medium or serum samples, the values of the sample channels were corrected for a “medium” or “albumin” channel, respectively, which reflects the unspecific and non-covalent adsorption of medium components or BSA/RSA to α-toxin and is generated by injection of incubation medium or 1% BSA/RSA in PBS, respectively, and further processing identical to the sample channels. To avoid the generation of air-bubbles (by spontaneous degassing, EDC/NHS reaction, pipetting or others), the buffers were degassed by applying vacuum (200 mbar for 30 min) and eventual air-bubbles removed immediately before injection (by gently tapping the vial). To avoid blockage of tubing, sterile-filtered buffers (0.2-μm sterile filters) and degassed distillated water (ddH_2_O) were used only and visually inspected for lack of particles or other contaminants. To avoid blockages in the system (autosampler- needle, autosampler, fluidic cell, tubing), it was cleaned after each experiment and on a regular basis weekly and monthly according to the instructions of the manufacturer.

### Enzymatic pretreatment of samples

#### Lipases

20 μl of serum or 4.5 ml of incubation medium were supplemented with 80 μl of 1x PI-PLC buffer (20 mM TRIS/HCl, pH 7.8, containing 0.1% BSA, 144 mM NaCl, 0.5 mM EDTA, 1 mM DTT and 0.1 mM PMSF), PLA_2_ buffer (25 mM HEPES/KOH, pH 7.2, containing 0.1% BSA, 144 mM NaCl, 2 mM Ca^2+^, 1 mM DTT and 0.1 mM PMSF) or PC-PLC buffer (20 mM TRIS/HCl, pH 7.8, containing 0.1% BSA, 144 mM NaCl, 2 mM Zn^2+^, 1 mM DTT and 0.1 mM PMSF) or GPI-PLD buffer (10 mM TRIS/maleate, pH 7.4, containing 144 mM NaCl, 1 mM Ca^2+^, 1 mM DTT and 0.1 mM PMSF) or 500 μl of the corresponding 10x buffers, respectively, and then incubated (30 min, 30°C) with partially purified PI-PLC from *Bacillus cereus* as described previously^39^ or PLA_2_ from honey bee venom or PC-PLC from *Chlostridium perfringens*^40^ or recombinant human GPI-PLD, respectively, at the concentrations indicated in the figure legends. <u>Proteinase K</u>: 50 μl of serum were supplemented with 200 μl of PBS and then incubated (60 min, 4°C) with proteinase K at the concentrations indicated in the absence or presence of 1 mM PMSF. *<u>N</u>*<u>-Glycanase F</u>: 50 μl of serum were supplemented with 200 μl of 0.1 M sodium phosphate buffer (pH 8.6), 1 mM DTT and 0.1 mM PMSF and then incubated (30 min, 30°C) with *N*-glycanase F from *Flavobacterium meningosepticum* at the concentrations indicated in the absence or presence of 0.5 M sodium thiocyanate. <u>α-</u> <u>Mannosidase</u>: 50 μl of serum were supplemented with 200 μl of 25 mM TRIS/HCl (pH 7.5) and 30 mM NaCl and then incubated (30 min, 30°C) with α-mannosidase from *Canavalla ensiformis* at the concentrations indicated in the absence or presence of 10 mM HgCl_2_.

### Chemical pretreatment of samples

Pretreatments for the chemical degradation of GPI structures were performed as described previously^41-44^ with the following modifications: <u>Dephosphorylation by hydrogen fluoride (HF)</u>: 50 μl of serum were supplemented with 200 μl of 50% (w/v) HF (T) or NaCl (C) and 0.5 M pyridine at −20°C in a polyethylene tube fitted with a cap. After incubation (60 min, 0 ± 0.5°C), the mixture was poured into a stirred saturated solution of LiOH at 4°C and then rapidly adjusted to pH 7.0 by the dropwise addition of 1 M LiOH. Prior to measurement, HF was removed under a stream of N_2_. <u>Deamination by nitrous acid (NA)</u>: 50 μl of serum were supplemented with 200 μl of 60 mM sodium acetate buffer (pH 4.0) containing 333 mM NaNO_3_ and then incubated (60 min, 22°C) as adapted from a previous report^41^.

### Immunoblotting

Proteins separated by sodium polyacrylamide gel electrophoresis (SDS-PAGE) were transferred to polyvinylidene difluoride membranes using the semi-dry procedure as described previously^12^. Washed membranes were incubated (24 h, 4°C) with anti-CD73 antibodies (1:1500), again washed and then incubated (3 h, 4°C) with horseradish peroxidase-coupled secondary goat anti-mouse IgG antibodies (1:4000). Labeled proteins were visualized by enhanced chemiluminescence. Lumiimages were evaluated with a luminescence imager using LumiImager software (Roche Diagnostics).

### Silver staining and densitometry

A protocol for “Stains-all” was used modified for enhanced sensitivity as described previously^45^. After SDS-PAGE (Novex precast gels, 10% acrylamide, BIS/TRIS resolving gel, morpholinopropanesulfonic acid-SDS running buffer), the gel was incubated (30 min) ten times with 100 ml of 25% isopropanol, each, on a shaker. Thereafter, the gel was incubated (25°C, 4-6 h) with 50 ml of “Stains-all” solution (30 mM TRIS/HCl, pH 8.7, 7.5% formamide, 25% isopropanol containing 0.025% [w/v] “Stains-all”) in light-tight containers on an orbital shaker. Subsequently, the gel was washed three times with 50 ml of 25% isopropanol, each, with a change every 10 min until the background became clear, then rinsed five times with water and finally incubated (40 min) in 50 ml of 12 mM silver nitrate on an orbital shaker. After rinsing of the gel with water and then with developer solution (0.28 M sodium carbonate and 0.15% formaldehyde) three times for 30 sec, each, the gel was incubated (5-10 min) with developer solution until the protein bands were visible. After termination of the developing reaction by removal of developer and the addition of 10% acetic acid, the gel was stored in a solution of 5% glycerol and 10% acetic acid. Densitometric scanning was performed using the Gel Print System (BioPhotonic Corp.) with the Video Copy Processor P4OU (Mitsubishi Inc.). Quantitative evaluation was performed utilizing the software package GPTools v3.0, 1-D Gel Analysis Software (BioPhotonic Corp.).

### Miscellaneous

Protein concentration was determined using the bicinchoninic acid protein assay kit (Thermo Fisher Scientific, Waltham, USA) with BSA as calibration standard.

### Statistics

Original data were analyzed and fitted using the FitMaster^®^ Origin-based software (Origin Inc.) upon subtraction of the corresponding values obtained with a blank and/or control channel as indicated. Statistical significance was determined with unpaired two-tailed t-test using GraphPad Prism 6 (version 6.0.2, GraphPad Software Inc.) software.

**Data availability statement.** Source data for **Figure 3b-h** and **Supplementary Fig. 10 and 11** are available online.

